# Structure of HIV-2 Nef reveals unique features distinct from HIV-1 involved in immune regulation

**DOI:** 10.1101/779439

**Authors:** Kengo Hirao, Sophie Andrews, Kimiko Kuroki, Hiroki Kusaka, Takashi Tadokoro, Shunsuke Kita, Toyoyuki Ose, Sarah L. Rowland Jones, Katsumi Maenaka

**Author notes:** These authors contributed equally: Kengo Hirao, Sophie Andrews, Kimiko Kuroki. Correspondence (K.M.), (S.R-J).

## Abstract

The HIV accessory protein Nef plays a major role in establishing and maintaining infection, particularly through immune evasion. Many HIV-2 infected people experience long-term viral control and survival, resembling HIV-1 elite control. HIV-2 Nef has overlapping but also distinct functions from HIV-1 Nef. Here we report the crystal structure of HIV-2 Nef core. The dileucine sorting motif forms a helix bound to neighboring molecules, and moreover, isothermal titration calorimetry demonstrated that the CD3 endocytosis motif can directly bind to HIV-2 Nef, ensuring AP-2 mediated endocytosis for CD3. The highly-conserved C-terminal region forms a α-helix, absent from HIV-1. We further determined the structure of SIV Nef harboring this region, demonstrating similar C-terminal α-helix, which may contribute to AP-1 binding for MHC-I downregulation. These results provide new insights into the distinct pathogenesis of HIV-2 infection.

## Introduction

Human Immunodeficiency Virus type 1 (HIV-1) is the main causative agent of Acquired Immunodeficiency Syndrome (AIDS). Whilst HIV-1 has spread throughout the world over a few decades, the other human AIDS virus, HIV-2, is largely confined to West Africa (de Silva et al., 2008), and appears to be steadily declining in prevalence (de Silva et al., 2008). HIV-1 and HIV-2 arose as a result of cross-species transmission of distinct simian immunodeficiency virus (SIV): HIV-1 from SIV infecting chimpanzees and gorillas in central Africa, and HIV-2 from SIV infecting sooty mangabeys in West Africa (Chen et al., 1997). There are striking differences in the natural history of infection between HIV-1 and HIV-2. The great majority of HIV-1-infected patients will progress to AIDS and death if not treated with anti-retroviral therapy (ART), but a small fraction (0.15%) maintain viral loads below detection and normal CD4+ T-cell counts, termed elite controllers or long-term non-progressors (Okulicz et al., 2009). In contrast, a much larger group of HIV-2 infected subjects (37% of a community HIV-2 cohort in Guinea-Bissau) experience long-term viral control, which appears to be clinically stable, unlike HIV-1 infection (van der Loeff et al., 2010). Nevertheless, most HIV-2 infected subjects will progress to AIDS without treatment, with a clinical picture indistinguishable from HIV-1 (Schim van der Loeff et al., 2002).

The differences in the natural history of infection with HIV-1 and HIV-2 could result from host or viral factors or, more likely, from a combination of the two. Whilst there are clear immune correlates of HIV-2 non-progression, such as the gag-specific CD8+ T-cell response (de Silva et al., 2013), other studies suggest that specific viral factors could play an important role (Onyango et al., 2010). HIV-2 elicits a potent broadly neutralizing antibody response in most infected people, leading to speculation that HIV-2 envelope might be one of the key factors accounting for the reduced pathogenicity of HIV-2. However, the structure of HIV-2 gp120 was recently determined, and showed remarkable similarities to HIV-1 gp120 (Davenport et al., 2016). This led us to consider other viral factors that may explain the clinical differences between HIV-1 and HIV-2. The retroviral accessory protein Nef is a potential contributing factor, since it is relevant in HIV and SIV replication and pathogenesis and functional differences between HIV-1 and HIV-2 Nef have been reported (Munch et al., 2005). Deletions in Nef are clearly linked with low viral load and delayed disease progression in both primates and humans (Deacon et al., 1995; Kestler et al., 1991; Kirchhoff et al., 1995; Oelrichs et al., 1998). Despite its small size, HIV-1 (206 residues), HIV-2 (256 residues), and SIVmac (263 residues), Nef displays a plethora of functions, including down-regulation of membrane proteins such as CD3, CD4, CD28 and MHC class I and II molecules by mediating sorting systems with phosphofurin acidic cluster sorting protein 1 and 2 (PACS-1 and −2) (Bell et al., 1998; Garcia and Miller, 1991; Schindler et al., 2003; Schwartz et al., 1996; Stumptner-Cuvelette et al., 2001; Swigut et al., 2001) and the adaptor proteins 1 and 2 (AP-1 and −2) complexes, up-regulation of the invariant chain (Ii) associated with immature MHC class II complexes (Schindler et al., 2003; Stumptner-Cuvelette et al., 2001), up-regulation of the Fas protein on infected cells (Xu et al., 1997), modulation of signaling pathways through interactions with kinases (Lang et al., 1997), and increasing the infectivity of virus particles by counteracting the host restriction factors SERINC3/5 (Rosa et al., 2015; Usami et al., 2015).

Whilst many of these functions are broadly conserved among HIV-1, HIV-2, and SIV Nefs, there are some differences in Nef functions among retrovirus families. For example, most HIV-2 and SIV Nefs can downregulate the T-cell receptor (TCR)-CD3 complex by clathrin adaptor protein complex 2 (AP-2)-mediated endocytosis, but HIV-1 and its precursor SIV (SIVcpz and SIVgor) Nefs have lost this ability: this is thought to be a key factor promoting the chronic immune activation observed in HIV-1 infection (Schindler et al., 2006). HIV-2 and most SIV Nefs downregulate the co-stimulatory molecule CD28 much more efficiently than those of HIV-1 and its precursors (Munch et al., 2005; Schindler et al., 2006). Furthermore, while HIV-1 Nef binds to the SH3 domain of some kinases, such as Fyn, Lck and Hck, with high affinity, HIV-2 and SIVmac Nef do not possess this ability, but are still able to bind to full-length Hck (Collette et al., 2000; Greenway et al., 1999; Lang et al., 1997). In addition to these functional differences, HIV-2 Nef and most SIV Nefs appear to employ different mechanisms from HIV-1 Nef to mediate the same functions. For example, HIV-2 and SIVmac Nef use their C-terminal region, which is absent in HIV-1 Nef, to downregulate MHC class I molecules by binding to the AP-1 complex (Munch et al., 2005; Swigut et al., 2000). Similarly, HIV-1 and HIV-2 Nefs have been shown to use overlapping but distinct domains to up-regulate Ii (Munch et al., 2005).

Whilst substantial progress has been made in understanding the structural aspects of HIV-1 and SIV Nefs (Alvarado et al., 2014; Jia et al., 2012; Kim et al., 2010; Ren et al., 2014), no structures of HIV-2 Nef were resolved to date that might help to explain the key differences between HIV-1 and HIV-2 function. Here, we present the crystal structure of the core region of HIV-2 Nef, based on a primary viral sequence derived from an HIV-2-infected subject from the Caio HIV-2 community cohort, Guinea-Bissau. HIV-2 Nef possesses a unique C-terminal α-helix that is present in HIV-2 and SIV Nefs but absent in HIV-1 (Kim et al., 2010; Manrique et al., 2017). Only one structure of SIV Nef construct harboring this C-terminal region was recently determined as a complex with AP-2 by cryo electron microscopy (EM), showing essentially a very similar C-terminal helix (Buffalo et al., 2019). Here we also prepared and crystallized the core domain of SIVmac Nef, including the C-terminal region. The crystal structure of SIVmac Nef also showed a C-terminal helix.

## Results

### Crystal structures of HIV-2 Nef

HIV-2 Nef proteins (wild-type and C193Y mutant) were expressed in *Escherichia coli* as soluble proteins and purified to show a single peak by size exclusion chromatography as a monomer (**Figure S1A**). HIV-2 Nef C193Y mutant was crystallized as described in Materials and Methods and diffracted to 2.07 Å (**Figure S1B,C**). Of note, the C193Y mutation on HIV-2 Nef did not alter the overall structure in solution, confirmed by CD spectra (**Figure S1D**). The structure of HIV-2 Nef protein was solved by molecular replacement using the HIV-1 structure (PDB: 1AVV) and refined to the final model with good stereochemistry (Table 1). The core structure of HIV-2 Nef consists of five α-helices (α2, α3, α5, α6, α7) and two β-strands (β1, β2) (Figur 1A,C). Comparison with the structures of HIV-1 Nef (1AVV (Arold et al., 1997), sequence identity 46%) and SIVmac Nef (3IK5 (Kim et al., 2010), sequence identity 70%) resulted in root mean square deviation (RMSD) values of 0.674 Å and 0.580 Å, respectively, demonstrating that the overall structure of HIV-2 Nef is almost identical to those of both the HIV-1 and SIVmac Nefs (Kim et al., 2010; Lee et al., 1996) (Figure 1B-D). The electron density of the N-terminal region (residues 90-103) and part of the central loop (residues 182-185 and 199-202) of HIV-2 Nef was disordered, as previously reported in HIV-1 and SIVmac Nef structures (Arold et al., 1997; Kim et al., 2010). However, unlike most of existing Nef crystal structures, part of the central loop was resolved and forms an α-helix (α4). This was visualized because its di-leucine motif (ExxxLϕ) EANYLL interacts with the hydrophobic crevice formed by α2 and α3 of a neighboring Nef molecule, stabilizing the otherwise flexible loop (**Figure S2A**). This helix structure of the central loop has been observed in some other Nef structures, where interactions with either the adaptor protein 2 (AP-2) or the Nef protein itself stabilize the complex (Horenkamp et al., 2011; Manrique et al., 2017; Ren et al., 2014).

**Table. 1.** Date collection and refinement statistics.

| 466 Statistic | HIV-2 Nef | SIVmac Nef |
| --- | --- | --- |
| 467 Data collection |  |  |
| 468 X-ray source | PF BL-5A | PF BL-17A |
| 469 Resolution (Å) | 50.0-2.07 | 41.6-2.00 |
| 470 | (2.11-2.07) | (2.05-2.00) |
| 471 Wave length (Å) | 1.0000 | 0.9800 |
| 472 Space group | *P* 2<sub>1</sub>2<sub>1</sub>2<sub>1</sub> | *P* 6<sub>5</sub>22 |
| 473 Cell dimensions |  |  |
| 474 a, b, c (Å) | a = 43.57, | a = 72.53, |
| 475 | b = 44.77, | b = 72.53, |
| 476 | c = 103.63 | c = 111.09 |
| 477 Total reflections | 90611 | 229843 |
| 478 Unique reflections | 12958 | 12304 |
| 479 | (619) | (884) |
| 480 *I* / $\sigma$ (*I*) | 23.7 (2.9) | 20 (3.2) |
| 481 Redundancy | 7.0 (6.4) | 18.7 (18.9) |
| 482 Completeness (%) | 100.0 (99.7) | 100 (100) |
| 483 *R*<sub>merge</sub> | 0.06 (0.53) | 0.11 (1.01) |
| 484 |  |  |
| 485 Refinement |  |  |
| 486 *R*<sub>work</sub> (%) | 20.5 | 19.5 |
| 487 *R*<sub>free</sub> (%) | 24.0 | 24.1 |
| 488 No. of protein residues | 141 | 154 |
| 489 RMSD bonds (Å) | 0.0023 | 0.0049 |
| 490 RMSD angles (°) | 0.55 | 0.55 |
| 491 Ramachandran |  |  |
| 492 favored (%) | 98.6 | 99.3 |
| 493 allowed (%) | 1.4 | 0.7 |
| 494 outlier (%) | 0 | 0 |
| 495 Average B factor (Å<sup>2</sup>) | 43.7 | 32.6 |

**Table. 1 Data collection and refinement statistics**
| 762 | Statistic | HIV-2 Nef | SIVmac Nef |
| --- | --- | --- | --- |
| 763 | Data collection |  |  |
| 764 | X-ray source | PF BL-5A | PF BL-17A |
| 765 | Resolution (Å) | 50.0-2.07 | 41.6-2.00 |
| 766 |  | (2.11-2.07) | (2.05-2.00) |
| 767 | Wave length (Å) | 1.0000 | 0.9800 |
| 768 | Space group | *P* 2<sub>1</sub>2<sub>1</sub>2<sub>1</sub> | *P* 6<sub>5</sub>22 |
| 769 | Cell dimensions |  |  |
| 770 | a, b, c (Å) | a = 43.57, | a = 72.53, |
| 771 |  | b = 44.77, | b = 72.53, |
| 772 |  | c = 103.63 | c = 111.09 |
| 773 | Total reflections | 90611 | 229843 |
| 774 | Unique reflections | 12958 | 12304 |
| 775 |  | (619) | (884) |
| 776 | *I* / $\sigma$ (*I*) | 23.7 (2.9) | 20 (3.2) |
| 777 | Redundancy | 7.0 (6.4) | 18.7 (18.9) |
| 778 | Completeness (%) | 100.0 (99.7) | 100 (100) |
| 779 | *R*<sub>merge</sub> | 0.06 (0.53) | 0.11 (1.01) |
| 780 |  |  |  |
| 781 | Refinement |  |  |
| 782 | *R*<sub>work</sub> (%) | 20.5 | 19.5 |
| 783 | *R*<sub>free</sub> (%) | 24.0 | 24.1 |
| 784 | No. of protein residues | 141 | 154 |
| 785 | RMSD bonds (Å) | 0.0023 | 0.0049 |
| 786 | RMSD angles (°) | 0.55 | 0.55 |
| 787 | Ramachandran |  |  |
| 788 | favored (%) | 98.6 | 99.3 |
| 789 | allowed (%) | 1.4 | 0.7 |
| 790 | outlier (%) | 0 | 0 |
| 791 | Average B factor (Å<sup>2</sup>) | 43.7 | 32.6 |
| 792 |  |  |  |

**Figure 1.**
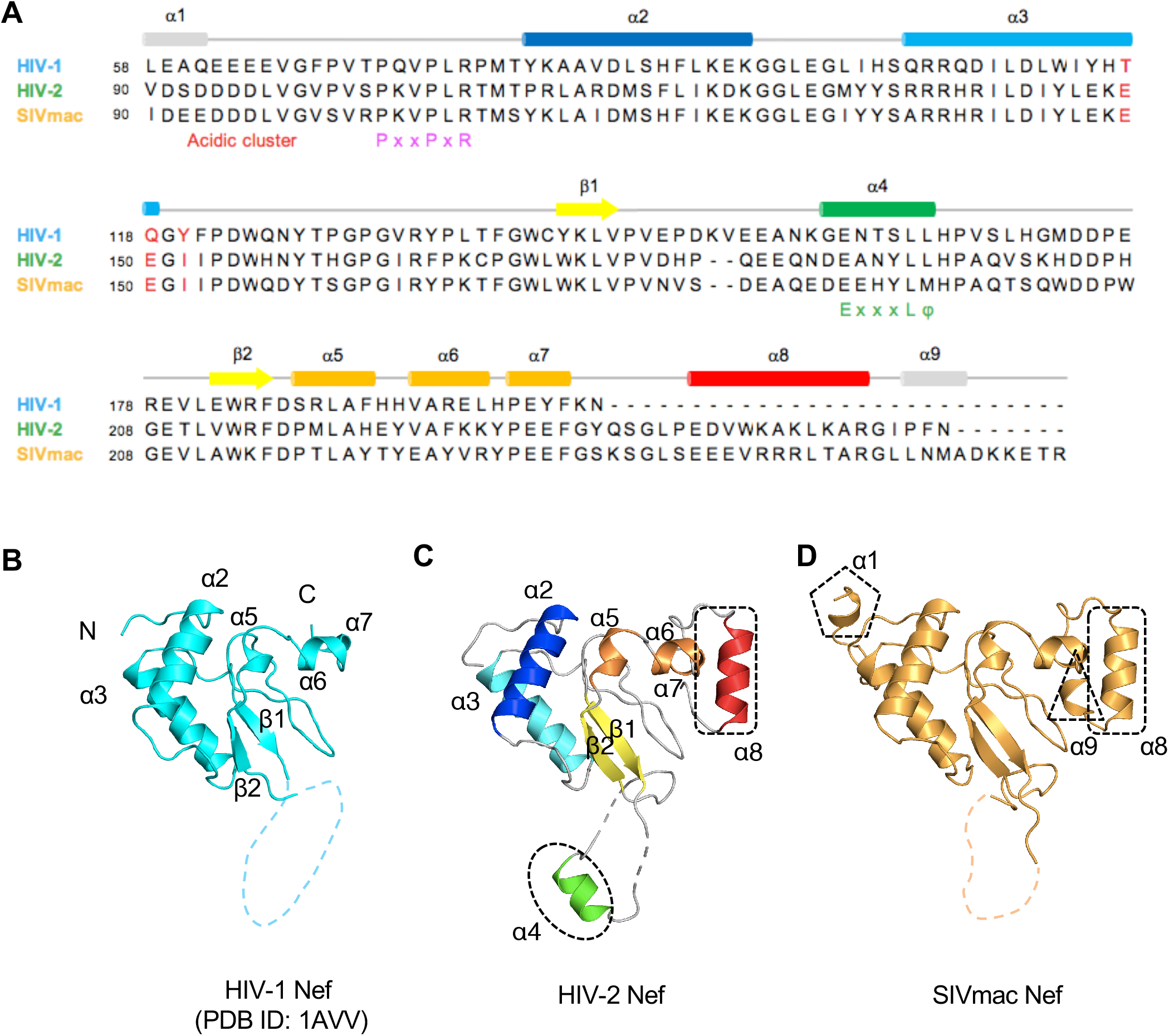
Crystal structures of HIV-1, HIV-2, and SIVmac Nef proteins. (A) Alignment of the Nef sequences of HIV-1, HIV-2, and SIVmac Nefs. The rods and arrows above the sequences indicate α-helix and β-sheet, respectively. The α-helix and β-sheet shown in blue mean these secondary structures are common structure among HIV-1, HIV-2, and SIVmac Nefs. (A) Structure of HIV-1 Nef (PDB ID: 1AVV) (C) Structure of HIV-2 Nef. (D) Structure of SIVmac239 Nef. (B ∼ D) Each structure is shown in Ribbon-model from the same orientation. Dotted circles indicate unique structures determined in HIV-2 and SIVmac Nefs.

### HIV-2 Nef contains a unique and conserved C-terminal alpha helix

A unique C-terminal α-helix (α8) was observed in HIV-2 Nef (Figure 1c **dotted square**). This structure is wholly absent in the HIV-1 protein. Ser237 in the loop between α7 and α8 helices forms a hydrogen bond with the main chain amine group of Leu239 to make an ST turn (Figure 2A **and Figure S3A**). This ST turn is often seen at the N-terminus of α-helices as a helix cap (Doig et al., 1997; Wan and Milner-White, 1999). Glu241 forms a hydrogen bond with Tyr235 and the highly-conserved Lys245 (Figure 2A **and Figure S3B**). The interaction between side chains of charged residues 3 to 4 positions apart, introducing charged residues on an adjacent turn of the α-helix, seems to increase the helix propensity. Arg251 makes a hydrogen bond network with His161, Glu231, and Glu232 to fix the α-helix in its place (Figure 2A **and Figure S3C**). Finally, Trp244 and Leu248 of the C-terminal helix are centrally located in a hydrophobic environment consisting of Glu231, Glu232, Tyr235, Glu241, Lys245, Ile253, and Phe255, making this helix fixed in the core structure (Figure 2A **and Figure S3D**). Fifty three HIV-2 group A Nef sequences obtained from the Los Alamos National Laboratory (LANL) HIV Sequence Database were analyzed by WebLogo (Crooks et al., 2004) (Figure 2C), revealing that many residues are conserved in this C-terminal region. A helical-wheel display of the C-terminal helix demonstrates that the highly conserved hydrophobic residues Trp244 and Leu248, together with Glu241 and Arg251, are located at the interface with the core structure (Figure 2D). In addition, the ST turn residues are also highly conserved. Taken together, these results suggest that the C-terminal structure is a conserved feature of HIV-2 Nef.

**Figure 2.**
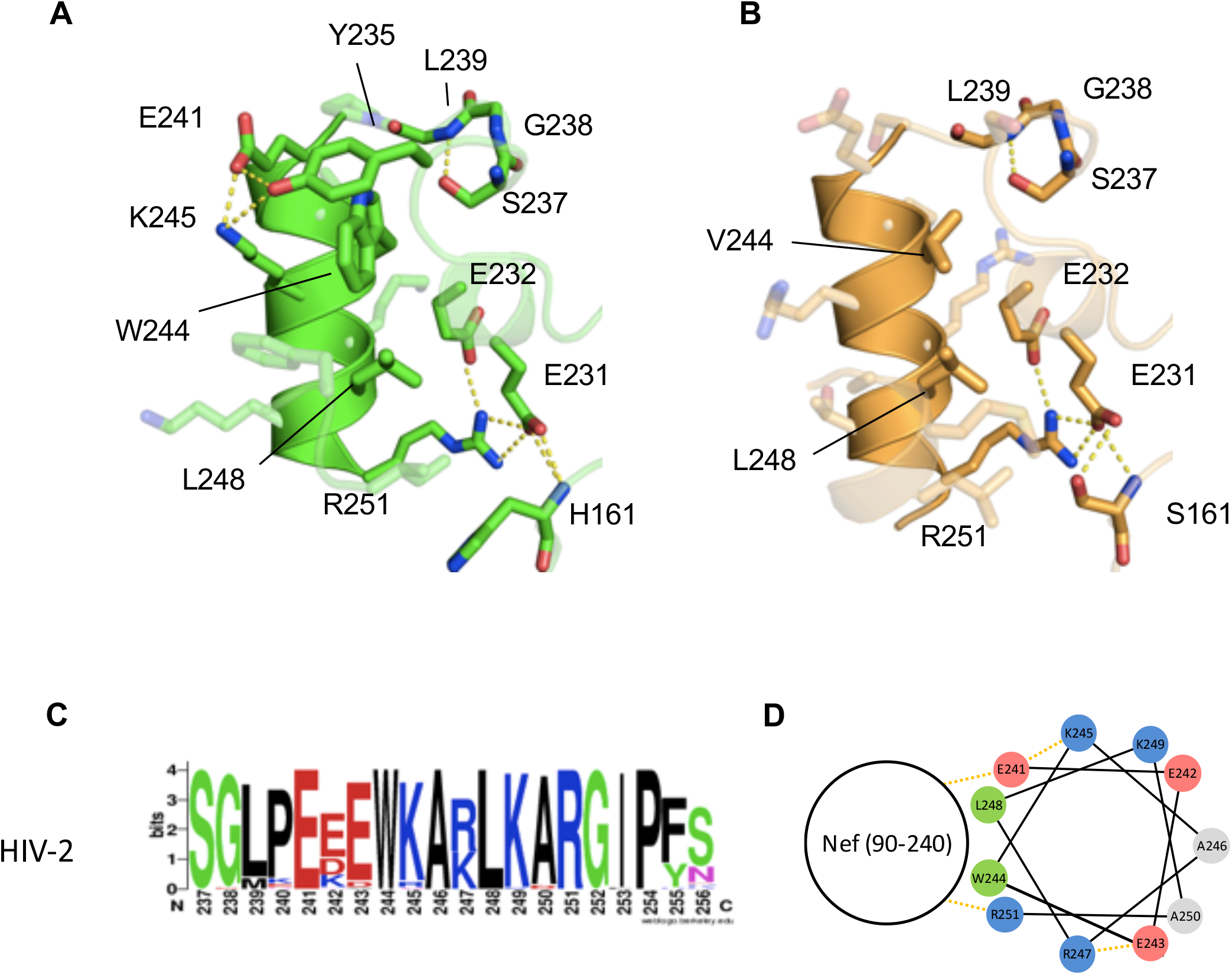
C-terminal structures of HIV-2 and SIVmac Nefs. (A and B) Important residues for the C-terminal helix structure in HIV-2 and SIVmac Nefs. In each figure, residues involved are shown in stick model. The dotted-line in yellow indicates a hydrogen-bond. (C) Amino acid conservation of the C-terminal residues of HIV-2. Sequences of HIV-2 A from Los Alamos HIV database were aligned and the conservation of each C-terminal residue was analyzed by WebLogo. Residue number for HIV-2 was based on Nef from present study (D) Helical-wheel display of the consensus sequence of HIV-2 Nef C-terminal α-helix (residue: 241-251)

### SIVmac Nef also has the C-terminal helix

Amino acid sequence alignment indicates that key residues of α8 helix of HIV2 Nef are essentially conserved with SIVmac Nef (Figure 1A). Whilst several crystal structures of SIVmac Nef core-domain structures have previously been resolved, the C-terminal region was removed prior to crystallization in all cases (Kim et al., 2010; Manrique et al., 2017). On the other hand, the cryo EM structure of SIVmac Nef complexed with AP-2 recently reported that it has a C-terminal helix (some residues between residues 235 and 250) (Buffalo et al., 2019). Here, the core domain of SIVmac Nef protein harboring the C-terminal region was prepared in a similar way to HIV-2 Nef (**Figure S1E**). The SIVmac Nef crystals diffracted to 2.0 Å (**Figure S1F,G**). The structure was determined by molecular replacement using HIV-2 Nef as a search model (Table 1).

The SIVmac Nef core structure is essentially the same as the reported truncated version of SIVmac Nef and similar to HIV2 Nef (RMS deviation of 0.43 Å for an overlay of 92 Cα atoms, Figure 1C,D**, Figure S2C**). The C-terminal α-helix (α8) was found in the present “full core” SIVmac Nef similarly to HIV2 Nef (Figure 1D **dotted square**). SIVmac Nef has more residues in the C-terminal region than HIV-2 Nef and the additional residues form a short α-helix (α9) (Figure 1D **dotted triangle and Figure S3E**). While detail description and PDB data is not currently available, similar helix formation was also presented in the recent paper of cryo EM structure of its AP-2 complex (Buffalo et al., 2019). The ST turn, the hydrogen bond network, and Val244 and Leu248 buried in a hydrophobic environment (Figure 2B **and Figure S3F-H**) closely resemble the HIV-2 Nef structure. These strongly conserved features suggest the functional significance of this novel structure, not only in HIV-2 but also in SIVmac Nefs.

### HIV-2 Nef interacts with di-leucine sorting motif in the common binding mode

EANYLL (residues 190-195), the di-leucine motif of HIV-2 Nef, interacts with the hydrophobic crevice formed by the α2 and α3 helices, as shown in Figure 3 **and Figure S2A**. This di-leucine motif is flexible but forms an α-helix (α4) when bound to the hydrophobic crevice, enabling two critical residues, Glu190 and Leu194, to align on the same side (Figure 3A). Glu190 interacts with a highly conserved Arg137 residue (Figure 3A). On the other hand, the two leucine residues of the motif, Leu194 and Leu195, fit into the hydrophobic crevice as follows (Figure 3B): Leu194 binds to the center of the crevice contacting Leu129, Met132, Arg138, Ile141, and Leu142, and the second leucine of the motif, Leu195, is deeply accommodated into the upper side of the crevice consisting of Ile123, Leu129, Leu142, Tyr145, and Leu146. Notably, a recent report of the SIVmac Nef structure (Manrique et al., 2017) showed that the dileucine sorting motif binds to the crevice formed by the α2 and α3 helices, similar to the current HIV-2 Nef structure (Figure 3C **and Figure S2C**), presumably reflecting the same sorting mechanism for CD3 and CD4 downregulation of HIV2 and SIVs.

**Figure 3.**
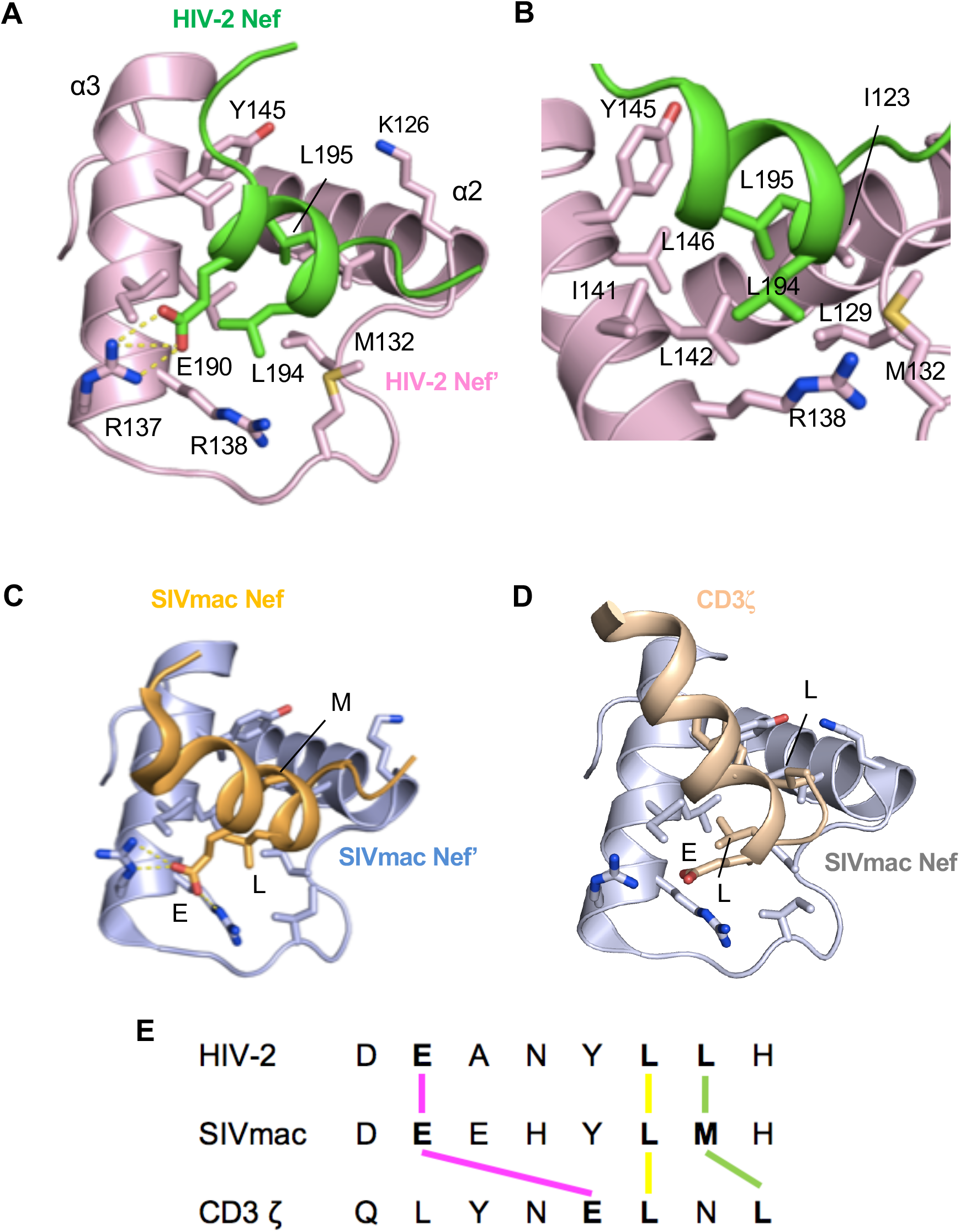
Common binding of sorting motifs. (A-D) Key residues involved in the interaction are shown in stick model. Each structure (A, C and D) is displayed from the same angle, while that of (B) is close-up view of (A) from different angle. (A and B) Structure of di-leucine motif (green) and the hydrophobic crevice of HIV-2 Nef. (C) Structure of di-leucine motif (dark-green) and the hydrophobic crevice of SIVmac239 Nef (light brown) (PDB: 5NUI). (D) Structure of CD3 zeta chain (light-green) and the hydrophobic crevice of SIVmac239 Nef (light brown) (PDB: 3IK5). (D) Display of the key binding sites of hydrophobic crevice with sorting motif. Glutamic acid residue interacts with R137 and R138 (pink). First leucine with hydrophobic residues, mainly with the residue at position 132 (yellow). Second leucine with hydrophobic residues, mainly with the residue at position 123 (green). The forth binding site (orange) binds to the corresponding residue for each motif. (E) Alignment of the sorting motifs.

### HIV-2 Nef has the capacity to interact with the CD3 zeta chain

The previously reported crystal structure of SIVmac complexed with the sorting motif of CD3 zeta chain further suggested that the binding mode of the motif is similar to those of the di-leucine motifs in HIV-2 and SIV Nefs (Figure 3D **and Figure S2D**) (Kim et al., 2010). Although the helical structure of CD3 zeta chain in SIVmac is tilted compared to that of the di-leucine motifs of HIV-2 and SIVmac Nefs, critical residues are rearranged so as to be located in the similar positions for the counterpart residues of the motifs (Figure 3D **and Figure S4**). Amino acid sequences of the motifs are different (Figure 3E) but they form an α-helix so that key glutamic acids interact with arginine residue(s) and two hydrophobic residues fit into the hydrophobic crevice (Figure 3). This raises the possibility that HIV-2 Nef has the ability to bind to the CD3 zeta chain in a similar way to SIVmac Nef. Indeed, HIV-2 Nef was previously reported to interact with CD3 zeta chain at a cellular level (Schaefer et al., 2002), however, the molecular basis for this interaction was poorly understood. Here, the binding of HIV-2 Nef with the CD3 zeta chain was investigated by isothermal titration calorimetry (ITC) measurements. The consensus HIV-1 and HIV-2 Nef proteins were prepared by the same method as HIV2 Nef (Figure 4A **and Figure S5A**), and a peptide was synthesized based on the previous studies, encompassing the second ITAM motif of CD3 zeta chain, termed SNID2. The ITC experiment showed that the dissociation constant of SIVmac Nef with the SNID2 peptide was 5.0 μM, which is in good agreement with the previous result (Manrique et al., 2017) (Figure 4D). As expected, no binding was observed for HIV-1 Nef (Figure 4B). On the other hand, HIV-2 Nef interacts with the peptide with a dissociation constant of 12.0 μM (Figure 4C). This result provides the first direct evidence that HIV-2 Nef can interact with the CD3 zeta chain without any additional factor *in vitro*.

**Figure 4.**
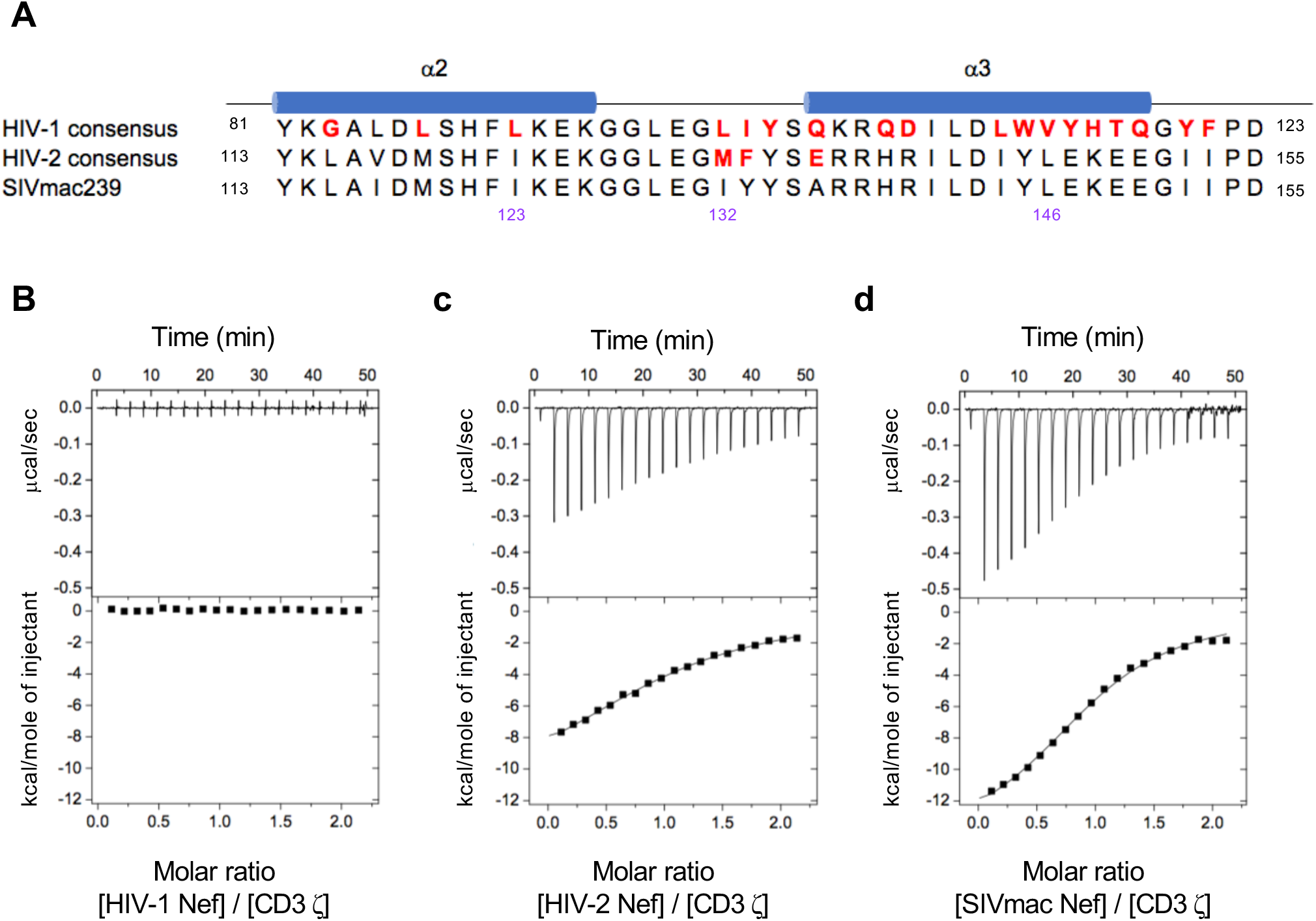
Interactions of Nef with CD3ζ. (A) Amino acid sequence alignment of the residues involved in CD3 ζ binding. (B) No binding was observed for HIV-1 consensus Nef. (C) HIV-2 consensus Nef interacted with CD3 ζ with a dissociation constant of 12.0 μM. (D) SIVmac239 Nef interacted with CD3 ζ with the highest affinity 5.0 μM.

Previously, a I132T mutation on HIV-2 Nef was shown to abrogate its ability to downregulate CD3 (Khalid et al., 2012). Here we introduced the same mutation to the best binding Nef, SIVmac Nef, to evaluate its effects on CD3 zeta chain binding. The SIVmac I132T Nef mutant showed modest heat generation but the binding constant could not be determined (**Figure S5B**). This suggests that Nef indeed interacts with the CD3 zeta chain using its hydrophobic crevice. In addition to the I132T mutation, a double mutation (I123L/L146F) has been also reported to disrupt the interaction of Nef with the CD3 zeta chain, resulting in loss of CD3 downregulation (Schmokel et al., 2013). The ITC experiment demonstrated that this mutation greatly disrupts the ability of Nef to interact with the CD3 zeta chain, even though slight residual binding tendency was retained (**Figure S5C**). Of note, both I132T and I123L/L146F mutations do not cause any overall structural changes, revealed by CD spectra (**Figure S5D,E**). Together with the functional data on I132T and I123L/L146F mutations, these suggest that Nef needs to interact with the CD3 zeta chain with a dissociation constant in the μM range to achieve CD3 downregulation.

### Structural comparison of Src-family kinase binding sites on Nefs explains distinct affinities

HIV-2 and SIVmac Nefs lack the ability to bind to the SH3 domain of Hck, one of the strongest interaction partners for HIV-1 Nef (Greenway et al., 1999; Karn et al., 1998; Lang et al., 1997). On the other hand, SIVmac Nef can bind to the SH3 domains of Fyn and Src, while HIV-1 Nef can also bind to them but more weakly than Hck (Collette et al., 2000). HIV-2 and SIVmac Nefs exhibit different preferences for binding SH3 domains from HIV-1. The proline-rich motif (PxxPxR) on HIV-1 Nef is essential for binding to the SH3 domain of Src family kinases, and is conserved among HIV-1, HIV-2, and SIVmac Nefs (Figure 1A). The main differences lie in residues Thr121, Gln122, and Tyr124 in HIV-1 Nef, which are represented by Glu149, Glu150, and Ile152, respectively, in HIV-2 and SIVmac sequences (Figure 1A). Among these residues, Tyr124 has been shown to play a critical role in the binding to the SH3 domain of Src kinases in a previous mutational study (Choi and Smithgall, 2004). Tyr124 in HIV-1 Nef directly interacts with Arg81, making the conformation of PxxPxR motif suitable for recognition by Trp114 of the SH3 domain, which is conserved in all Src family kinases (Figure 5A). On the other hand, Tyr124 also forms a hydrophobic pocket together with Met83, contributing to make hydrophobic interaction with Ile92 of the SH3 domain. In contrast, the equivalent residue in HIV-2 and SIVmac, Ile152, does not seem capable of contributing sufficiently in either fashion (Figure 5C,E). This critical amino acid difference accounts for the weak binding of HIV-2 and SIVmac Nefs to SH3 domains of Src family kinases.

**Figure 5.**
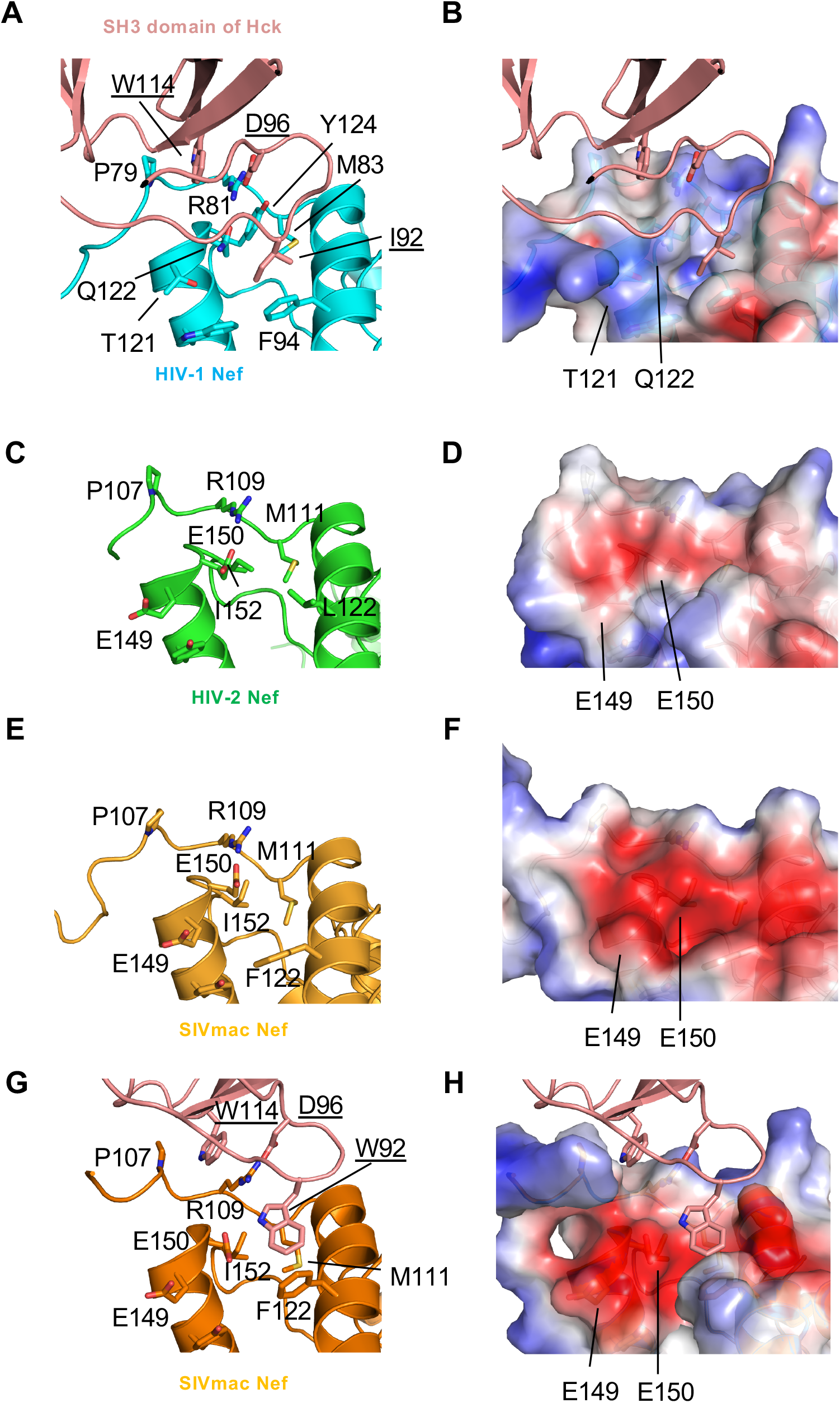
Distinct Src-family kinase binding affinity. (A) HIV-1 Nef (PDB ID: 4U5W) complexed with the SH3 domain of Hck is shown in Ribbon model. (B) HIV-1 Nef complexed with Hck (PDB ID: 4U5W). The surface of HIV-1 Nef is colored based on the calculated potentials from −3.0 kT/e (red) to +3.0 kT/e (blue). (C, E, G) Zoomed-in view in HIV-2 Nef (C), SIVmac Nef (E), SIVmac Nef complexed with HCK I92W mutant (PDB ID: 5NUH) (G). (D, F, H) Electrostatic potential representation of HIV-2 Nef (D), SIVmac Nef (F) and SIVmac Nef complexed with HCK I92W mutant (H).

The electrostatic aspects of the HIV-1 Nef structure demonstrated a positively-charged surface for SH3 domain binding (Figure 5B). In contrast, this surface in HIV-2 and SIVmac Nefs is negatively-charged because of the presence of Glu149 and Glu150 residues (Figure 5D,F). This electrostatic presentation might explain why HIV-2 and SIVmac Nefs show a preference for binding to Fyn and Src kinases, which have a positively charged residue at position 92. On the other hand, the SIVmac Nef was successfully crystallized as its complex with the I92W mutant of HCK kinase, which has higher affinity, (Manrique et al., 2017), showing some different recognition mode (Figure 5G,H). Trp92 of HCK faces vertically to Phe122 of Nef, pushing the surrounding residues far from Glu149 and Glu150 (Nef), while a salt bridge between Asp96 (HCK) and Arg81 (Nef) is maintained. These amino acid differences between HIV-1 and HIV-2/SIV Nefs have an effect on the preference of the SH3 domain binding, but it is also worth noting that that the distinct pattern of the amino acid differences overlaps with the residues responsible for CD3 binding, suggesting that while HIV-2 and SIVmac Nef do not possess the capacity to bind strongly to the SH3 domain of Src family kinases, they retained the ability to interact with the CD3 zeta chain instead.

### N-terminal acidic cluster interacts with the hydrophobic crevice in SIVmac Nef

In the present study, the structure of the N-terminal acidic cluster (residues 91 - 96, corresponding to residues 59 – 64 in HIV-1) of SIVmac Nef was determined (Figure 1D**, dotted pentagon and** Figure 6). The N-terminal region includes an acidic cluster and interacts with the hydrophobic crevice formed by the α2 and α3 helices of a neighboring Nef molecule, which is utilized for binding to the di-leucine motif in HIV-2 Nef (**Figure S2B**), while the central loop of SIVmac Nef (residues 180-210) containing the motif was disordered. Interestingly, N-terminal residues are extended in a direction toward the α2 - but not the α3 - helix (Figure 6A,B). Upon binding to the hydrophobic crevice, a part of the acidic cluster forms a short α-helix, allowing Ile90 and Asp94 to align in the same direction (Figure 6A). The previous report of crystal structure of HIV-1Nef complexed with its protein inhibitor demonstrated that the N-terminal acidic cluster forms a longer helix with Trp57, preceding the N-terminal acidic cluster, accommodating into the hydrophobic pocket (Breuer et al. 2011) (Figure 6C). This recognition mode is somehow distinct from that of shorter helix with extended conformation in the present structure of SIVmac Nef (Figure 6B), which harbors aspartic acid at the site corresponding to Trp57 (this construct does not include this residue). Notably, this binding in the current structure comprises of more extended interaction with six acidic residues in the current structure and is very similar to those of sorting motifs observed in Figure 3; 1) Asp94 forms a salt bridge with Arg137 and 2) Ile90 fits into the hydrophobic crevice consisting of Ile123, Gly128, Leu129, Arg138, Ile141, Leu142. This could be a common feature explaining how Nef accommodates its binding partner into the hydrophobic crevice. In addition to these interactions, other acidic residues (Asp91, Glu92, Glu93, Asp95 and Asp96) also contribute to the binding, mainly by contacting the positively charged surface (Figure 6B). The recent cryo-EM structure of HIV-1 Nef complexed with AP-1 and the crystal structure of the complex of AP-1 μ subunit revealed that acidic cluster forms an extended conformation to interact widely with the positively charged surface of the adjacent μ subunit (Figure 6D) (Jia et al., 2012; Morris et al., 2018). The present structure, however, showed another conformation of this region of Nef, implying that it may assume multiple conformations depending on the situation.

**Figure 6.**
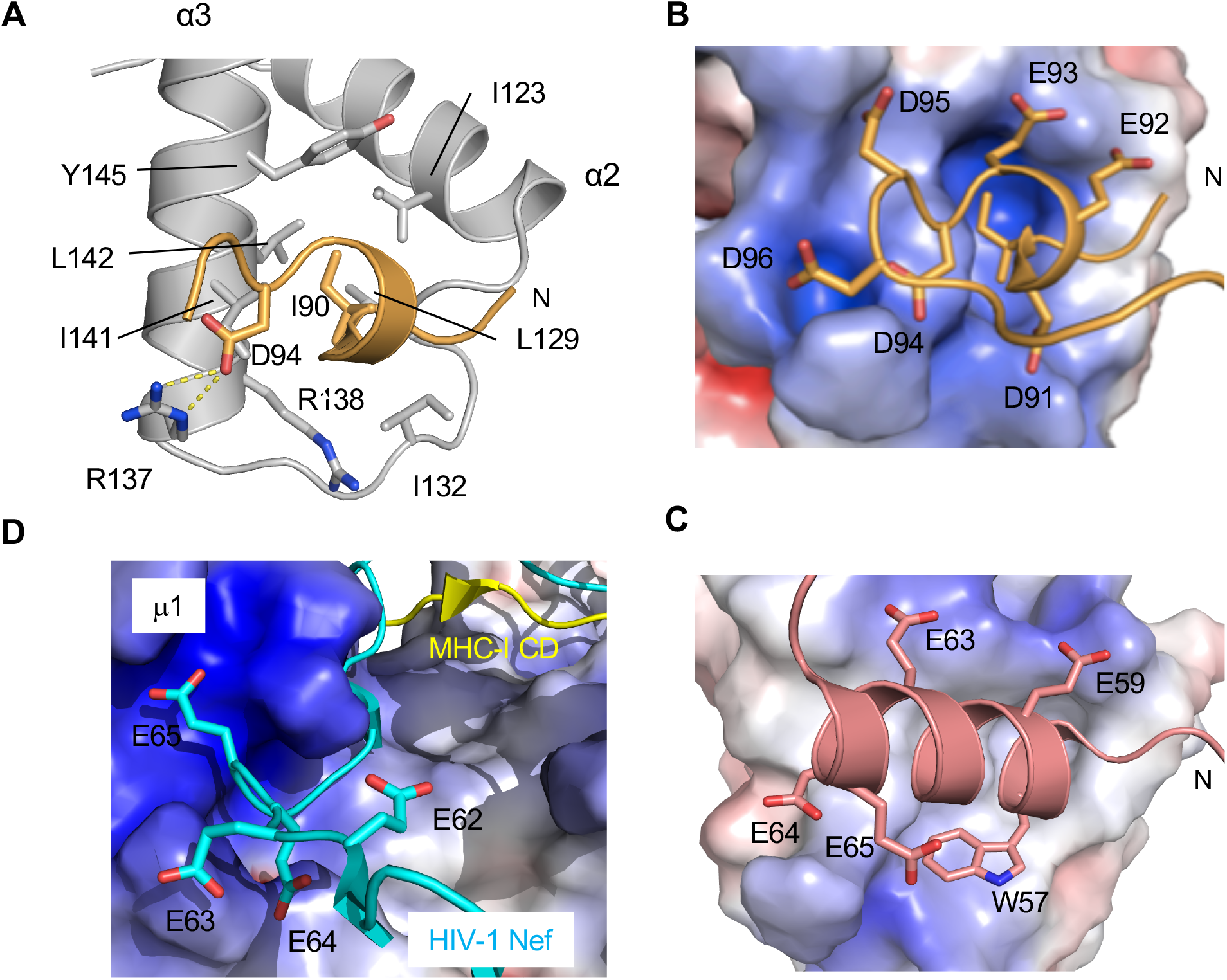
Structural plasticity of the N-terminal acid cluster. (A) Binding of the acidic-cluster (orange) with the hydrophobic crevice of neighboring Nef. Side chains of residues involved in the interaction are shown in stick model and labeled. (B) Binding of the acidic cluster (orange) with the hydrophobic crevice of neighboring Nef in surface model. Side chains of residues are shown in stick model. The surface of SIVmac Nef is colored based on the calculated potentials from −5.0 kT/e (red) to +5.0 kT/e (blue). (C) Binding of the acidic cluster (salmon pink) with the hydrophobic crevice in HIV1 in complex with engineered Hck-SH3 domain (PDB ID: 3REA). The surface of HIV1 Nef is colored as same as (B). (D) Different conformation of acidic-cluster of Nef. Acidic cluster of HIV-1 Nef (cyan) are shown in stick model and it forms extended conformation along with the positive charged surface of μ subunit (PDB ID: 4EN2). The surface of μ subunit is colored as same as (B).

## Discussion

Here we present the first crystal structures of HIV-2 and SIVmac Nefs to demonstrate their “full” core regions. The overall structure of HIV-2 Nef is essentially similar to both those of HIV-1 and SIVmac, emphasizing the importance of their core structures. The present HIV-2 Nef structure also revealed how its di-leucine motif binds to a hydrophobic crevice formed by the α2 and α3 helices, providing the binding modes conserved with SIV Nefs. Furthermore, a unique C-terminal helix was found in both HIV-2 and SIVmac Nefs and is essentially conserved between HIV-2 and SIVmac Nefs, implying its functional significance.

Comparison of the present structure with that of SIVmac Nef bound to the CD3 zeta chain gave us important insights into how HIV-2 Nef interacts with the motif of the CD3 zeta chain. Three critical residues, Glu190 and two hydrophobic residues, Leu194 and Leu195, in the di-leucine motif of Nef and Glu74, Leu75, and Leu77 in CD3 zeta chain, are located at the same binding sites for each residue despite their different sequential order (Figure 3). The glutamic acid residues, Glu190 in Nef and Glu74 in the CD3 zeta chain, strongly interact with well-conserved arginine residues Arg137 and Arg138. The previous mutagenesis study revealed that these residues are relevant for the ability of HIV-2 and SIVmac Nefs to downmodulate CD3 (Manrique et al., 2017). Two hydrophobic residues of the di-leucine motif LL in HIV-2 and CD3 zeta chain (**<u>L</u>**N**<u>L</u>**) fit into the deep crevice. As expected by structural analysis together with the previous study on CD3 zeta chain binding (Manrique et al., 2017), the ITC experiment here showed that HIV-2 Nef binds the CD3 zeta chain with a similar dissociation constant to SIVmac Nef. The I132T or I123L/L146F mutations greatly reduce the affinity, emphasizing the importance of the intricate features of the crevice. This result also indicates that direct interaction between Nef and CD3 zeta chain with μM order of dissociation constant is required for CD3 down-regulation.

Despite its overall similarity, the present structures provide insight into the underlying reasons for the differential affinities to the SH3 domain of HIV-1, HIV-2, and SIVmac Nefs. While the proline-rich motif and other residues involved in the interaction with the SH3 domain are well conserved, Thr117, Gln118, and Tyr120 in HIV-1 Nef are systematically represented by Glu149, Glu150, and Ile152 in HIV-2 and SIVmac, respectively. These different residues completely change the characteristics of the binding sites for the SH3 domain, leading to the loss of the ability to bind to the SH3 domain of Hck and different preferences for Fyn and Src kinases. However, SIVmac Nef has evolved to interact with Hck via a different mechanism (Greenway et al., 1999). Its close relative HIV-2 Nef, whose putative SH3-binding site structurally resembles that of SIVmac, likely uses a similar binding mechanism. The fact that Nefs target the same kinases in different ways emphasizes their importance for the virus life-cycle *in vivo*.

The present SIVmac Nef structure revealed the structure of the flexible N-terminal acidic cluster by interacting with the hydrophobic crevice as observed in Figure 3A. The structure formed an α-helix so that Ile90 and Asp94 can be aligned to make important interactions (Figure 6A). Furthermore, other acidic residues strengthen the interaction with the positively charged surface of the crevice (Figure 6B). The way that this hydrophobic crevice accommodates the motifs share a lot in common and the crevice has been used to interact with various binding partners, for example binding the cytoplasmic tail of CD4. The di-leucine motif of the CD4 cytoplasmic tail is necessary for its downregulation by HIV-1, HIV-2, and SIV Nefs (Hua and Cullen, 1997). Likewise, the di-leucine motif of CD28 is essential for its downregulation (Swigut et al., 2001). As observed in the binding with CD3 zeta chain, this hydrophobic crevice might play a role in binding to the cytoplasmic tail of CD28 and result in the different potencies of its downregulation by HIV-1 and HIV-2/SIVmac Nefs. Furthermore, although the N-terminal region interacts with the crevice of its neighboring Nef, the N-terminus orients towards the α2 helix and this binding can happen intermolecularly. Indeed, a previous NMR structural study showed that the flexible N-terminus binds to its crevice in solution (Grzesiek et al., 1997). Of note, the acidic cluster has been implicated in PACS binding and the hydrophobic crevice was recently suggested as the likely binding site (Dikeakos et al., 2012). Therefore, Nef might accommodate the N-terminal acidic cluster until it finds its correct binding partner such as PACS. This model was proposed some time ago, and the present structure might help our understanding of how Nef accommodates its flexible N-terminus and hides the multi-binding “hub”, the hydrophobic crevice.

In this study, a unique C-terminal structural element (α8) is revealed. The residues at the α8 helix and its surrounding region are well conserved not only in HIV-2 but also in SIVmac Nefs, suggesting a strong evolutionary pressure, implying functional significance. The importance of the additional C-terminus of Nef was previously demonstrated by an investigation showing that a SIVmac strain harboring C-terminally truncated Nef was able to recover its virulence by obtaining additional C-terminal residues, comparable to that of SIVmac239 Nef (Lafont et al., 2000). It has also previously been shown that deletion of the C-terminal residues of HIV-2 and SIVmac239 Nef specifically inhibit their ability to downregulate MHC-I without abrogating other functions. MHC-I downregulation by HIV-2 and SIVmac Nefs is not as dependent on their proline-rich motifs as is HIV-1 Nef (Munch et al., 2005; Swigut et al., 2000). The present study allows us to provide a possible explanation for this difference. Firstly, the down-regulation of MHC-I involves the activation of Src-family kinases, however, activation is not dependent on the proline-rich motif of HIV-2 and SIVmac Nefs. Therefore, this step does not seem to be critical for the difference. On the other hand, the crystal structure of HIV-1 Nef in complex with the MHC-I cytoplasmic tail and the μ1 subunit of the adaptor protein 1 (AP-1) demonstrates extensive interactions of the proline-rich motif of HIV-1 Nef with the MHC-I tail. The recent cryo-EM structures of the HIV-1 Nef/AP-1 complex and HDX-MS analysis indicate that these tight interactions contribute to the open conformer of AP-1/HIV-1 Nef/MHC-I complex trimers, fitting to the clathrin pathway. When HIV-2 and SIVmac Nefs are superimposed on the HIV-1 Nef/MHC-I tail/μ1 subunit complex, the unique C-terminal structure can be seen to encapsulate the μ1 subunit without any steric hindrance (Figure 7**, Figure S6**). The C-terminal residues of HIV-2 and SIVmac Nefs could therefore provide an additional binding site to strengthen the AP-1 complex formation. This result raises the possibility that the C-terminal region tightly and/or suitably binds to AP-1 complex to induce an open conformer suitable for entry into the clathrin pathway.

**Figure 7.**
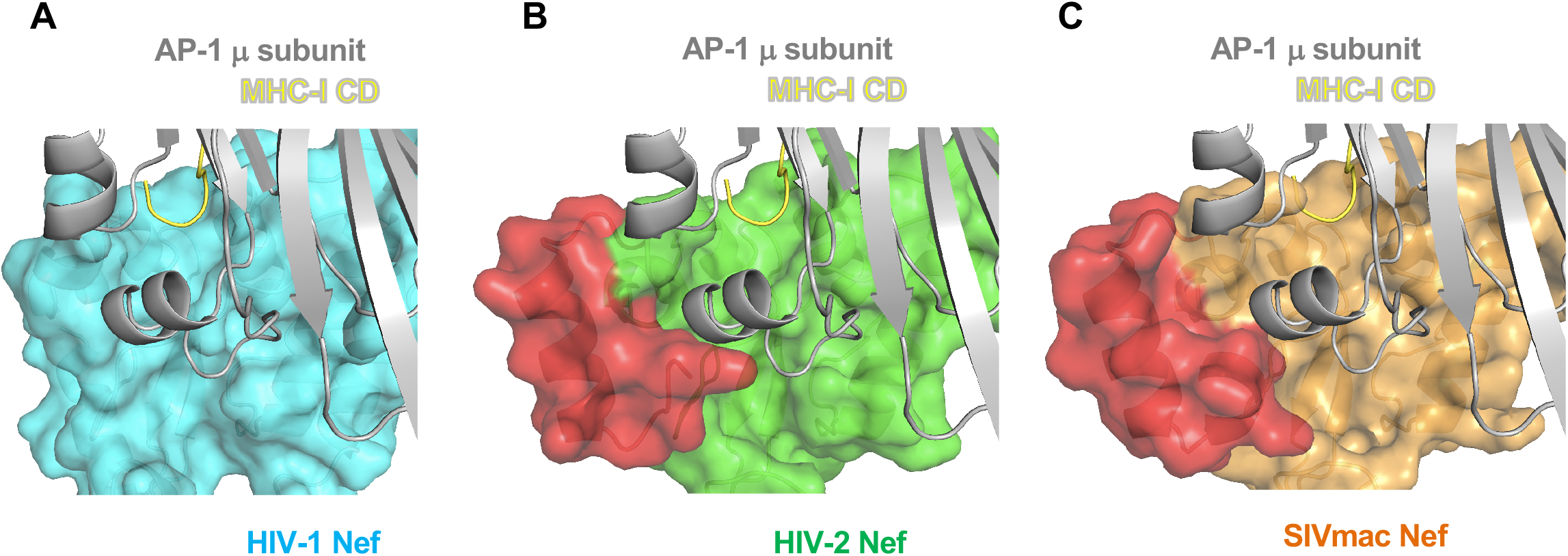
Models of AP-1 complexes with HIV-2 and SIVmac Nefs based on crystal structure of HIV-1 Nef complexed with AP-1 μ subunit and cytoplasmic tail of MHC class I molecule. (A) HIV-1 Nef (cyan) in complex with MHC-I CD (yellow) and the mu1 subunit of AP-1 (dark blue) (PDB 4EN2). (B) (C) Superimposition of HIV-2 Nef (B) and SIVmac Nef (C) onto MHC-I CD and μ1 subunit of AP1.

The present study provides another insight into the functional differences at a molecular level between HIV-1, HIV-2, and SIVmac Nefs. HIV-2 and SIVmac Nefs share the similar features including the unique C-terminal structure, unexpectedly discovered here, to play important roles in virus replication and immune evasion. Several differences, however, are still known between HIV-2 and SIVmac Nefs, such as the requirement for the CD4 tail for its down-modulation (Hua and Cullen, 1997). Future studies of the neglected N-terminal and C-terminal regions of Nefs, exhibiting relatively poor conservation among different lentiviruses, will further reveal the precise mechanisms of immune evasion, which in turn should significantly contribute to the design of effective anti-HIV therapies and vaccines.

## Acknowledgements

We thank the beam-line staff of Photon Factory (Tsukuba, Japan) for technical help during data collection. This work is supported in part by the Japan Society for the Promotion of Science (JSPS) Grants-in-Aid for Scientific Research KAKENHI (Grants 22121007), JSPS Strategic Young Researcher Overseas Visits Program for Accelerating Brain Circulation, Platform Project for Supporting Drug Discovery and Life Science Research (Basis for Supporting Innovative Drug Discovery and Life Science Research (BINDS)) from AMED under Grant Number 18am0101093j0002, and the Middle Molecule Project from AMED under Grant Number 18ae0101047h, Hokkaido University, the Global Facility Center (GFC), the Pharma Science Open Unit (PSOU), funded by the Ministry of Education, Science, Sports, Culture and Technology under the ‘‘Support Program for Implementation of New Equipment Sharing System’’, the Ministry of Health and, Labor and Welfare of Japan, Hokkaido University Biosurface project and Takeda Science Foundation.

## Author Contributions

K.H., S.A., K.K, S.R-J and K.M. designed research; K.H., S.A., K.K, H.K., and K.M. performed research; K.H., H.K., T.T., T.O., S.R-J and K.M. analyzed the data; K.H., K.K, S.R-J and K.M. wrote the paper.

## STAR Methods

### Plasmids

The full-length *nef* gene was isolated from the genomic (proviral) DNA of an HIV-2-infected donor (TD062) from the Caió community cohort, Guinea-Bissau. This subject was first diagnosed with HIV-2 infection in 1989, and by the time of sampling (2010) had progressed to a high plasma viral load (139, 519 copies/ml plasma) and CD4+ T-cell count of 497 cells/μL. In keeping with previous studies in this cohort, the sequence corresponded to HIV-2 clade A, which is the predominant subtype in this region of West Africa.

For protein expression, the full core domain (residues 90-256) was further amplified by PCR with primers containing NdeI and BamHI restriction sites. The resulting product was ligated into pET-16b (Novagen) between NdeI and BamHI of the multiple cloning site (pET-16b-HIV-2 Nef). To achieve crystallization of the molecule, a C193Y mutation was introduced using pET-16b-HIV-2 as a template by PCR-mediated site-directed mutagenesis (pET-16b-HIV-2 Nef C193Y).

The full-length *nef* gene of SIVmac239 was synthesized by eurofins. The premature stop codon (TAA) of SIVmac239 *nef* at 93^rd^ codon was changed to GAA coding glutamic acid as previously reported(Kestler et al., 1991). The full core domain (residues 90-263) was amplified and ligated (pET-16b-SIVmac239 Nef) as described above. Two mutants (I132T and I123L/L146F) of SIVmac239 Nef were prepared by PCR-mediated site-directed mutagenesis. For crystallization, pET-16b-SIVmac239 Nef was further modified with 3C protease cleavage site. (pET-16b-3C-SIVmac239 Nef)

The expression plasmids coding each core domain (HIV-1 consensus: residues 58-206, HIV-2 consensus: residues 90-256) were prepared as described above. Sequences of all constructs were confirmed by DNA sequences. The primers used in the present study were listed in **Supplementary Note**.

### Protein expression and purification

HIV-1, HIV-2, and SIVmac Nefs were expressed in *Escherichia coli* strain Rosetta2(DE3) as N-terminal His-tagged fusion proteins. The bacterial cells were grown in 2 x YT medium supplemented with 100 μg/mL ampicillin at 37°C to an optical density at 600 nm (OD_600_) of 0.6. Expression of the His-tagged proteins in the cells was induced with 0.1 mM isopropyl-β-D-thiogalactopyranoside (IPTG) at 16°C for 16 hours. The cells were harvested by centrifugation, re-suspended in 50 mM Tris-HCl (pH 8.0), 100 mM NaCl, and 10 mM imidazole, sonicated and cleared by centrifugation at 40,000 x g for 30 minutes. The supernatant was incubated with Ni-NTA agarose (QIAGEN) at 4°C for 2 hours with rotation. The mixture was washed, and the protein was eluted with 50 mM Tris-HCl (pH 8.0), 100 mM NaCl, and 200 mM imidazole. Elution fractions were pooled.

For crystallization of HIV-2 Nef, the buffer was exchanged into 20 mM Tris-HCl (pH 7.5), 100 mM NaCl, 2 mM CaCl_2_ by dialysis and the N-terminal His-tag was digested by FactorXa (QIAGEN) at 20°C overnight. For crystallization of SIVmac239 Nef, the buffer was exchanged into 50 mM Tris-HCl (pH 8.0), 100 mM NaCl, 5 mM dithiothreitol (DTT) by dialysis and the N-terminal His-tag was digested by 3C protease (kindly provided by) at 4°C overnight. The samples were further purified by size exclusion chromatography on a HiLoad 26/60 Superdex75 pg (GE healthcare) column in 50 mM Tris-HCl (pH 8.0), 100 mM NaCl, and 2 mM 2-Mercaptoethanol.

For ITC measurements, the elution fractions of Ni-NTA purification were directly subjected to size exclusion chromatography in 20 mM HEPES (pH 8.0), 100 mM NaCl, and 2 mM 2-Mercaptoethanol. Product purity was finally confirmed by SDS-PAGE.

The peptide of CD3 zeta chain containing SNID2 (residues: 115-QKDKMAEAYSEIGMKGERRRG-135) was synthesized by eurofins. The peptide was dissolved in 20 mM HEPES (pH 8.0), 100 mM NaCl for ITC measurements.

### Measurement of the CD spectra

CD spectra were measured on a Jasco J-820 spectropolarimeter (Japan Spectroscopic) using a scanning wavelength of 260-200 nm at 25°C. The proteins at 10 μM in 20 mM Tris-HCl (pH 8.0) was loaded into a quartz cell (optical path length: 1 mm). CD spectra of eight scans was averaged and the baseline was corrected by subtracting the averaged buffer spectra. Finally, ellipticity was converted to mean residue ellipticity.

### Crystallization of HIV-2 Nef and SIVmac239 Nef

The initial crystal of the HIV-2 Nef C193Y mutant was grown using the sitting-drop method at 293K by mixing 0.2 μL of the protein sample (10 mg/mL) with an equal volume of the reservoir solution containing 0.1 M MES (pH 6.5) and 30% polyethylene glycol (PEG) 300. The condition was optimized to 0.1 M MES (pH 6.5) and 10% PEG 3350 using the hanging-drop method at 277K by mixing 1 μL of the protein sample with an equal volume of the reservoir solution. The crystal was cryoprotected in the same reservoir solution and was flash-cooled in liquid nitrogen. The crystals of SIVmac239 were grown using the sitting-drop method at 293K by mixing 0.5 μL of the protein sample (12 mg/mL) with 1 μL of the reservoir solution containing 0.2 M Ammonium acetate, 0.1 M Sodium citrate (pH 5.6) and 30% PEG 4000. The crystal was cryoprotected consisting of reservoir solution supplemented with 12% glycerol and was flash-cooled in liquid nitrogen.

### Data collection and processing

Diffraction datasets of HIV-2 Nef C193Y and SIVmac239 Nef were collected on beamline BL-5A and BL-17A, respectively, at the Photon Factory, KEK, in Tsukuba, Japan. The datasets were processed with XDS (Kabsch, 2010). The structures of HIV-2 Nef and SIVmac239 Nef were determined by molecular replacement with HIV-1 Nef (Protein Data Bank [PDB] ID: 1AVV)(Arold et al., 1997) or solved HIV-2 Nef, respectively, using the program Molrep(Vagin A, 1997) implemented in the CCP4i suite(Winn et al., 2011). The structures were then refined using the program REFMAC(Murshudov et al., 2011) and Phenix(Adams et al., 2010) and manually fitted with COOT(Emsley et al., 2010). Root-mean-square deviation (RMSD) values of reported structures from our crystal structure were calculated using the “align” command in PyMOL version 1.8. The electrostatic potential of our structure was visualized with the APBS tools implemented in PyMOL version 1.8.

### Isothermal titration calorimetry

Interactions between various Nefs and the CD3 zeta chain were studied by Isothermal titration calorimetry (ITC) using MicroCal ITC200 microcalorimeter (GE Healthcare). Nef at concentration at 200 μM was stepwise injected from the syringe to 20 μM CD3 zeta chain place in the measurement cell at 25°C. The changes in heating power were monitored. Data were analyzed using the software provided by the manufacturer and the binding constant was fitted using one set of sites model.

### Protein Structure Accession Number

Atomic coordinates and structure factors for HIV-2 Nef and SIVmac239 Nef were deposited in the Protein Data Bank under the accession codes, 6K6M and 6K6N, respectively.

## References

Adams, P.D., Afonine, P.V., Bunkoczi, G., Chen, V.B., Davis, I.W., Echols, N., Headd, J.J., Hung, L.W., Kapral, G.J., Grosse-Kunstleve, R.W., et al. (2010). PHENIX: a comprehensive Python-based system for macromolecular structure solution. Acta Crystallogr D Biol Crystallogr 66, 213–221.

Alvarado, J.J., Tarafdar, S., Yeh, J.I., and Smithgall, T.E. (2014). Interaction with the Src homology (SH3-SH2) region of the Src-family kinase Hck structures the HIV-1 Nef dimer for kinase activation and effector recruitment. J Biol Chem 289, 28539–28553.

Arold, S., Franken, P., Strub, M.P., Hoh, F., Benichou, S., Benarous, R., and Dumas, C. (1997). The crystal structure of HIV-1 Nef protein bound to the Fyn kinase SH3 domain suggests a role for this complex in altered T cell receptor signaling. Structure 5, 1361–1372.

Bell, I., Ashman, C., Maughan, J., Hooker, E., Cook, F., and Reinhart, T.A. (1998). Association of simian immunodeficiency virus Nef with the T-cell receptor (TCR) zeta chain leads to TCR down-modulation. J Gen Virol 79 ( *Pt 11*), 2717–2727.

Breuer S, Schievink SI, Schulte A, Blankenfeldt W, Fackler OT, Geyer M. (2011). Molecular design, functional characterization and structural basis of a protein inhibitor against the HIV-1 pathogenicity factor Nef. PLoS One. 2011;6(5):e20033.

Buffalo, C.Z., Stürzel, C.M., Heusinger, E., Kmiec, D., Kirchhoff, F., Hurley, J.H., Ren, X. (2019) Structural basis for Tetherin antagonism as a barrier to zoonotic lentiviral transmission. Cell Host Microbe 26 (*3*), 359–368.

Chen, Z., Luckay, A., Sodora, D.L., Telfer, P., Reed, P., Gettie, A., Kanu, J.M., Sadek, R.F., Yee, J., Ho, D.D., et al. (1997). Human immunodeficiency virus type 2 (HIV-2) seroprevalence and characterization of a distinct HIV-2 genetic subtype from the natural range of simian immunodeficiency virus-infected sooty mangabeys. J Virol 71, 3953–3960.

Choi, H.J., and Smithgall, T.E. (2004). Conserved residues in the HIV-1 Nef hydrophobic pocket are essential for recruitment and activation of the Hck tyrosine kinase. J Mol Biol 343, 1255–1268.

Collette, Y., Arold, S., Picard, C., Janvier, K., Benichou, S., Benarous, R., Olive, D., and Dumas, C. (2000). HIV-2 and SIV nef proteins target different Src family SH3 domains than does HIV-1 Nef because of a triple amino acid substitution. J Biol Chem 275, 4171–4176.

Crooks, G.E., Hon, G., Chandonia, J.M., and Brenner, S.E. (2004). WebLogo: a sequence logo generator. Genome Res 14, 1188–1190.

Davenport, Y.W., West, A.P., Jr., and Bjorkman, P.J. (2016). Structure of an HIV-2 gp120 in Complex with CD4. J Virol 90, 2112–2118.

de Silva, T.I., Cotten, M., and Rowland-Jones, S.L. (2008). HIV-2: the forgotten AIDS virus. Trends Microbiol 16, 588–595.

de Silva, T.I., Peng, Y., Leligdowicz, A., Zaidi, I., Li, L., Griffin, H., Blais, M.E., Vincent, T., Saraiva, M., Yindom, L.M., et al. (2013). Correlates of T-cell-mediated viral control and phenotype of CD8(+) T cells in HIV-2, a naturally contained human retroviral infection. Blood 121, 4330–4339.

Deacon, N.J., Tsykin, A., Solomon, A., Smith, K., Ludford-Menting, M., Hooker, D.J., McPhee, D.A., Greenway, A.L., Ellett, A., Chatfield, C., et al. (1995). Genomic structure of an attenuated quasi species of HIV-1 from a blood transfusion donor and recipients. Science 270, 988–991.

Dikeakos, J.D., Thomas, L., Kwon, G., Elferich, J., Shinde, U., and Thomas, G. (2012). An interdomain binding site on HIV-1 Nef interacts with PACS-1 and PACS-2 on endosomes to down-regulate MHC-I. Mol Biol Cell 23, 2184–2197.

Doig, A.J., MacArthur, M.W., Stapley, B.J., and Thornton, J.M. (1997). Structures of N-termini of helices in proteins. Protein Sci 6, 147–155.

Emsley, P., Lohkamp, B., Scott, W.G., and Cowtan, K. (2010). Features and development of Coot. Acta Crystallogr D Biol Crystallogr 66, 486–501.

Garcia, J.V., and Miller, A.D. (1991). Serine phosphorylation-independent downregulation of cell-surface CD4 by nef. Nature 350, 508–511.

Greenway, A.L., Dutartre, H., Allen, K., McPhee, D.A., Olive, D., and Collette, Y. (1999). Simian immunodeficiency virus and human immunodeficiency virus type 1 nef proteins show distinct patterns and mechanisms of Src kinase activation. J Virol 73, 6152–6158.

Grzesiek, S., Bax, A., Hu, J.S., Kaufman, J., Palmer, I., Stahl, S.J., Tjandra, N., and Wingfield, P.T. (1997). Refined solution structure and backbone dynamics of HIV-1 Nef. Protein Sci 6, 1248–1263.

Horenkamp, F.A., Breuer, S., Schulte, A., Lulf, S., Weyand, M., Saksela, K., and Geyer, M. (2011). Conformation of the dileucine-based sorting motif in HIV-1 Nef revealed by intermolecular domain assembly. Traffic 12, 867–877.

Hua, J., and Cullen, B.R. (1997). Human immunodeficiency virus types 1 and 2 and simian immunodeficiency virus Nef use distinct but overlapping target sites for downregulation of cell surface CD4. J Virol 71, 6742–6748.

Jia, X., Singh, R., Homann, S., Yang, H., Guatelli, J., and Xiong, Y. (2012). Structural basis of evasion of cellular adaptive immunity by HIV-1 Nef. Nat Struct Mol Biol 19, 701–706.

Kabsch, W. (2010). Xds. Acta Crystallogr D Biol Crystallogr 66, 125–132.

Karn, T., Hock, B., Holtrich, U., Adamski, M., Strebhardt, K., and Rubsamen-Waigmann, H. (1998). Nef proteins of distinct HIV-1 or −2 isolates differ in their binding properties for HCK: isolation of a novel Nef binding factor with characteristics of an adaptor protein. Virology 246, 45–52.

Kestler, H.W., 3rd, Ringler, D.J., Mori, K., Panicali, D.L., Sehgal, P.K., Daniel, M.D., and Desrosiers, R.C. (1991). Importance of the nef gene for maintenance of high virus loads and for development of AIDS. Cell 65, 651–662.

Khalid, M., Yu, H., Sauter, D., Usmani, S.M., Schmokel, J., Feldman, J., Gruters, R.A., van der Ende, M.E., Geyer, M., Rowland-Jones, S., et al. (2012). Efficient Nef-mediated downmodulation of TCR-CD3 and CD28 is associated with high CD4+ T cell counts in viremic HIV-2 infection. J Virol 86, 4906–4920.

Kim, W.M., Sigalov, A.B., and Stern, L.J. (2010). Pseudo-merohedral twinning and noncrystallographic symmetry in orthorhombic crystals of SIVmac239 Nef core domain bound to different-length TCRzeta fragments. Acta Crystallogr D Biol Crystallogr 66, 163–175.

Kirchhoff, F., Greenough, T.C., Brettler, D.B., Sullivan, J.L., and Desrosiers, R.C. (1995). Brief report: absence of intact nef sequences in a long-term survivor with nonprogressive HIV-1 infection. N Engl J Med 332, 228–232.

Lafont, B.A., Riviere, Y., Gloeckler, L., Beyer, C., Hurtrel, B., Paule Kieny, M., Kirn, A., and Aubertin, A.M. (2000). Implication of the C-terminal domain of nef protein in the reversion to pathogenicity of attenuated SIVmacBK28-41 in macaques. Virology 266, 286–298.

Lang, S.M., Iafrate, A.J., Stahl-Hennig, C., Kuhn, E.M., Nisslein, T., Kaup, F.J., Haupt, M., Hunsmann, G., Skowronski, J., and Kirchhoff, F. (1997). Association of simian immunodeficiency virus Nef with cellular serine/threonine kinases is dispensable for the development of AIDS in rhesus macaques. Nat Med 3, 860–865.

Lee, C.H., Saksela, K., Mirza, U.A., Chait, B.T., and Kuriyan, J. (1996). Crystal structure of the conserved core of HIV-1 Nef complexed with a Src family SH3 domain. Cell 85, 931–942.

Manrique, S., Sauter, D., Horenkamp, F.A., Lulf, S., Yu, H., Hotter, D., Anand, K., Kirchhoff, F., and Geyer, M. (2017). Endocytic sorting motif interactions involved in Nef-mediated downmodulation of CD4 and CD3. Nat Commun 8, 442.

Morris, K.L., Buffalo, C.Z., Sturzel, C.M., Heusinger, E., Kirchhoff, F., Ren, X., and Hurley, J.H. (2018). HIV-1 Nefs Are Cargo-Sensitive AP-1 Trimerization Switches in Tetherin Downregulation. Cell 174, 659–671 e614.

Munch, J., Schindler, M., Wildum, S., Rucker, E., Bailer, N., Knoop, V., Novembre, F.J., and Kirchhoff, F. (2005). Primary sooty mangabey simian immunodeficiency virus and human immunodeficiency virus type 2 nef alleles modulate cell surface expression of various human receptors and enhance viral infectivity and replication. J Virol 79, 10547–10560.

Murshudov, G.N., Skubak, P., Lebedev, A.A., Pannu, N.S., Steiner, R.A., Nicholls, R.A., Winn, M.D., Long, F., and Vagin, A.A. (2011). REFMAC5 for the refinement of macromolecular crystal structures. Acta Crystallogr D Biol Crystallogr 67, 355–367.

Oelrichs, R., Tsykin, A., Rhodes, D., Solomon, A., Ellett, A., McPhee, D., and Deacon, N. (1998). Genomic sequence of HIV type 1 from four members of the Sydney Blood Bank Cohort of long-term nonprogressors. AIDS Res Hum Retroviruses 14, 811–814.

Okulicz, J.F., Marconi, V.C., Landrum, M.L., Wegner, S., Weintrob, A., Ganesan, A., Hale, B., Crum-Cianflone, N., Delmar, J., Barthel, V., et al. (2009). Clinical outcomes of elite controllers, viremic controllers, and long-term nonprogressors in the US Department of Defense HIV natural history study. J Infect Dis 200, 1714–1723.

Onyango, C.O., Leligdowicz, A., Yokoyama, M., Sato, H., Song, H., Nakayama, E.E., Shioda, T., de Silva, T., Townend, J., Jaye, A., et al. (2010). HIV-2 capsids distinguish high and low virus load patients in a West African community cohort. Vaccine 28 *Suppl 2*, B60–67.

Ren, X., Park, S.Y., Bonifacino, J.S., and Hurley, J.H. (2014). How HIV-1 Nef hijacks the AP-2 clathrin adaptor to downregulate CD4. Elife 3, e01754.

Rosa, A., Chande, A., Ziglio, S., De Sanctis, V., Bertorelli, R., Goh, S.L., McCauley, S.M., Nowosielska, A., Antonarakis, S.E., Luban, J., et al. (2015). HIV-1 Nef promotes infection by excluding SERINC5 from virion incorporation. Nature 526, 212–217.

Schaefer, T.M., Bell, I., Pfeifer, M.E., Ghosh, M., Trible, R.P., Fuller, C.L., Ashman, C., and Reinhart, T.A. (2002). The conserved process of TCR/CD3 complex down-modulation by SIV Nef is mediated by the central core, not endocytic motifs. Virology 302, 106–122.

Schim van der Loeff, M.F., Jaffar, S., Aveika, A.A., Sabally, S., Corrah, T., Harding, E., Alabi, A., Bayang, A., Ariyoshi, K., and Whittle, H.C. (2002). Mortality of HIV-1, HIV-2 and HIV-1/HIV-2 dually infected patients in a clinic-based cohort in The Gambia. AIDS 16, 1775–1783.

Schindler, M., Munch, J., Kutsch, O., Li, H., Santiago, M.L., Bibollet-Ruche, F., Muller-Trutwin, M.C., Novembre, F.J., Peeters, M., Courgnaud, V., et al. (2006). Nef-mediated suppression of T cell activation was lost in a lentiviral lineage that gave rise to HIV-1. Cell 125, 1055–1067.

Schindler, M., Wurfl, S., Benaroch, P., Greenough, T.C., Daniels, R., Easterbrook, P., Brenner, M., Munch, J., and Kirchhoff, F. (2003). Down-modulation of mature major histocompatibility complex class II and up-regulation of invariant chain cell surface expression are well-conserved functions of human and simian immunodeficiency virus nef alleles. J Virol 77, 10548–10556.

Schmokel, J., Li, H., Shabir, A., Yu, H., Geyer, M., Silvestri, G., Sodora, D.L., Hahn, B.H., and Kirchhoff, F. (2013). Link between primate lentiviral coreceptor usage and Nef function. Cell Rep 5, 997–1009.

Schwartz, O., Marechal, V., Le Gall, S., Lemonnier, F., and Heard, J.M. (1996). Endocytosis of major histocompatibility complex class I molecules is induced by the HIV-1 Nef protein. Nat Med 2, 338–342.

Stumptner-Cuvelette, P., Morchoisne, S., Dugast, M., Le Gall, S., Raposo, G., Schwartz, O., and Benaroch, P. (2001). HIV-1 Nef impairs MHC class II antigen presentation and surface expression. Proc Natl Acad Sci U S A 98, 12144–12149.

Swigut, T., Iafrate, A.J., Muench, J., Kirchhoff, F., and Skowronski, J. (2000). Simian and human immunodeficiency virus Nef proteins use different surfaces to downregulate class I major histocompatibility complex antigen expression. J Virol 74, 5691–5701.

Swigut, T., Shohdy, N., and Skowronski, J. (2001). Mechanism for down-regulation of CD28 by Nef. EMBO J 20, 1593–1604.

Usami, Y., Wu, Y., and Gottlinger, H.G. (2015). SERINC3 and SERINC5 restrict HIV-1 infectivity and are counteracted by Nef. Nature 526, 218–223.

Vagin A, T.A. (1997). MOLREP: an Automated Program for Molecular Replacement. J Appl Cryst 30, 1022–1025.

van der Loeff, M.F., Larke, N., Kaye, S., Berry, N., Ariyoshi, K., Alabi, A., van Tienen, C., Leligdowicz, A., Sarge-Njie, R., da Silva, Z., et al. (2010). Undetectable plasma viral load predicts normal survival in HIV-2-infected people in a West African village. Retrovirology 7, 46.

Wan, W.Y., and Milner-White, E.J. (1999). A recurring two-hydrogen-bond motif incorporating a serine or threonine residue is found both at alpha-helical N termini and in other situations. J Mol Biol 286, 1651–1662.

Winn, M.D., Ballard, C.C., Cowtan, K.D., Dodson, E.J., Emsley, P., Evans, P.R., Keegan, R.M., Krissinel, E.B., Leslie, A.G., McCoy, A., et al. (2011). Overview of the CCP4 suite and current developments. Acta Crystallogr D Biol Crystallogr 67, 235–242.

Xu, X.N., Screaton, G.R., Gotch, F.M., Dong, T., Tan, R., Almond, N., Walker, B., Stebbings, R., Kent, K., Nagata, S., et al. (1997). Evasion of cytotoxic T lymphocyte (CTL) responses by nef-dependent induction of Fas ligand (CD95L) expression on simian immunodeficiency virus-infected cells. J Exp Med 186, 7–16.

