## Supplemental Information for "Structure of HIV-2 Nef reveals unique features distinct from HIV-1 involved in immune regulation"

**Supplementary text**

**Primers used for generation of core domain of Nef**

HIV-1 Nef consensus

F:	5’-GACATATCATATGCTAGAAGCACAAGAGGAGGAGG-3’

R:	5’-GACATAGGATCCTCAGCAGTCCTTGTAGTACTCCA-3’

HIV-2 Nef consensus

F:	5’-GACATATCATATGGTAGATTCAGATGATGATGACC-3’

R:	5’-GACATAGGATCCTCATCAACTAAATGGTATCCCTC-3’

HIV-2 Nef WT

F:	5’-GACATATCATATGGTAGATTCAGATGATGATGACC-3’

R:	5’-GACATAGGATCCTCACTAATTAAATGGTATCCCTC-3’

SIVmac239 Nef

F:	5’-GACATATTCATATGATAGATGAGGAAGATGATGAC-3’

R:	5’-GACATAGGATCCTCATCAGCGAGTT TCCTTCTTGT-3’

**Primers used for introducing mutation**

HIV-2 Nef C193Y

F:	5’-GAGGCTAAC**TAC**TTACTGCA-3’

R:	5’-TGCAGTAA**GTA**GTTAGCCTC-3’

SIVmac239 Nef I132T

F:	5’-CTGGAAGGG**ACT**TATTACAG-3’

R:	5’-CTGTAATA**AGT**CCCTTCCAG-3’

SIVmac239 Nef I123L

F:	5’-TCTCATTTT**CTA**AAAGAAAA-3’

R:	5’-TTTTCTTT**TAG**AAAATGAGA-3’

SIVmac239 Nef L146F

F:	5’-GACATATAC**TTT**GAAAAGGA-3’

R:	5’-TCCTTTTC**AAA**GTATATGTC-3’

**Primers used for introducing 3C site**

3C site

F:	5’-TTCTGTTCCAGGGGCCCATAGATGAGGAAGATGAT-3’

R:	5’-TGGAACAGAACTTCCAGCATATGACGACCTTCGAT-3’

**Amino acid sequence used in this study**

>His-HIV-1 Nef consensus

MGHHHHHHHHHHSSGHIEGRHMLEAQEEEEVGFPVRPQVPLRPMTYKGALDLSHFLKEKGGLEGLIYSQKRQDILDLWVYHTQGYFPDWQNYTPGPGIRYPLTFGWCFKLVPVEPEKVEEANEGENNSLLHPMSQHGMDDPEKEVLVWKFDSRLAFHHMARELHPEYYKDC

>His-HIV-2 Nef consensus

MGHHHHHHHHHHSSGHIEGRHMVDSDDDDLVGVPVTPRVPLRAMTYKLAVDMSHFIKEKGGLEGMFYSERRHRILDIYLEKEEGIIPDWQNYTHGPGIRYPMFFGWLWKLVPVDVPQEGEDNETHCLMHPAQTSRFDDPHGETLVWRFDPMLAYEYKAFIRYPEEFGHKSGLPEEEWKARLKARGIPFS

>His-HIV-2 Nef WT

MGHHHHHHHHHHSSGHIEGRHMVDSDDDDLVGVPVSPKVPLRTMTPRLARDMSFLIKDKGGLEGMYYSRRRHRILDIYLEKEEGIIPDWHNYTHGPGIRFPKCPGWLWKLVPVDHPQEEQNDEANCLLHPAQVSKHDDPHGETLVWRFDPMLAHEYVAFKKYPEEFGYQSGLPEDVWKAKLKARGIPFN

>His-HIV-2 Nef C193Y

MGHHHHHHHHHHSSGHIEGRHMVDSDDDDLVGVPVSPKVPLRTMTPRLARDMSFLIKDKGGLEGMYYSRRRHRILDIYLEKEEGIIPDWHNYTHGPGIRFPKCPGWLWKLVPVDHPQEEQNDEANYLLHPAQVSKHDDPHGETLVWRFDPMLAHEYVAFKKYPEEFGYQSGLPEDVWKAKLKARGIPFN

>HM-HIV-2 Nef C193Y

HMVDSDDDDLVGVPVSPKVPLRTMTPRLARDMSFLIKDKGGLEGMYYSRRRHRILDIYLEKEEGIIPDWHNYTHGPGIRFPKCPGWLWKLVPVDHPQEEQNDEANYLLHPAQVSKHDDPHGETLVWRFDPMLAHEYVAFKKYPEEFGYQSGLPEDVWKAKLKARGIPFN

>His-SIVmac239 Nef

MGHHHHHHHHHHSSGHIEGRHMIDEEDDDLVGVSVRPKVPLRTMSYKLAIDMSHFIKEKGGLEGIYYSARRHRILDIYLEKEEGIIPDWQDYTSGPGIRYPKTFGWLWKLVPVNVSDEAQEDEEHYLMHPAQTSQWDDPWGEVLAWKFDPTLAYTYEAYVRYPEEFGSKSGLSEEEVRRRLTARGLLNMADKKETR

>His-SIVmac239 Nef I132T

MGHHHHHHHHHHSSGHIEGRHMIDEEDDDLVGVSVRPKVPLRTMSYKLAIDMSHFIKEKGGLEGTYYSARRHRILDIYLEKEEGIIPDWQDYTSGPGIRYPKTFGWLWKLVPVNVSDEAQEDEEHYLMHPAQTSQWDDPWGEVLAWKFDPTLAYTYEAYVRYPEEFGSKSGLSEEEVRRRLTARGLLNMADKKETR

>His-SIVmac239 Nef I123L/L146F

MGHHHHHHHHHHSSGHIEGRHMIDEEDDDLVGVSVRPKVPLRTMSYKLAIDMSHFLKEKGGLEGIYYSARRHRILDIYFEKEEGIIPDWQDYTSGPGIRYPKTFGWLWKLVPVNVSDEAQEDEEHYLMHPAQTSQWDDPWGEVLAWKFDPTLAYTYEAYVRYPEEFGSKSGLSEEEVRRRLTARGLLNMADKKETR

>His-**3C**-SIVmac239 Nef

MGHHHHHHHHHHSSGHIEGRHM**LEVLFQGP**IDEEDDDLVGVSVRPKVPLRTMSYKLAIDMSHFIKEKGGLEGIYYSARRHRILDIYLEKEEGIIPDWQDYTSGPGIRYPKTFGWLWKLVPVNVSDEAQEDEEHYLMHPAQTSQWDDPWGEVLAWKFDPTLAYTYEAYVRYPEEFGSKSGLSEEEVRRRLTARGLLNMADKKETR

>GP-SIVmac239 Nef

GPIDEEDDDLVGVSVRPKVPLRTMSYKLAIDMSHFIKEKGGLEGIYYSARRHRILDIYLEKEEGIIPDWQDYTSGPGIRYPKTFGWLWKLVPVNVSDEAQEDEEHYLMHPAQTSQWDDPWGEVLAWKFDPTLAYTYEAYVRYPEEFGSKSGLSEEEVRRRLTARGLLNMADKKETR

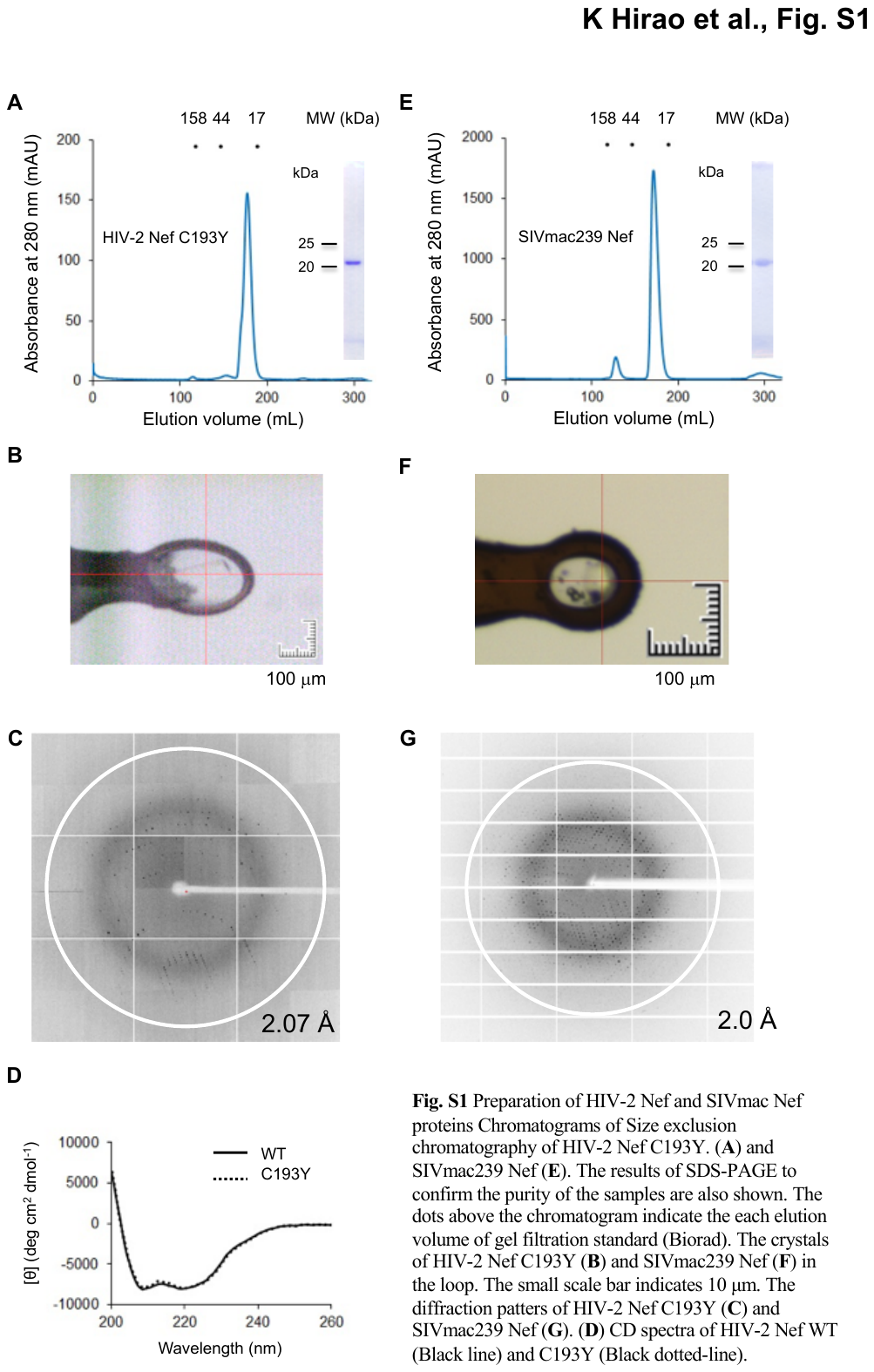

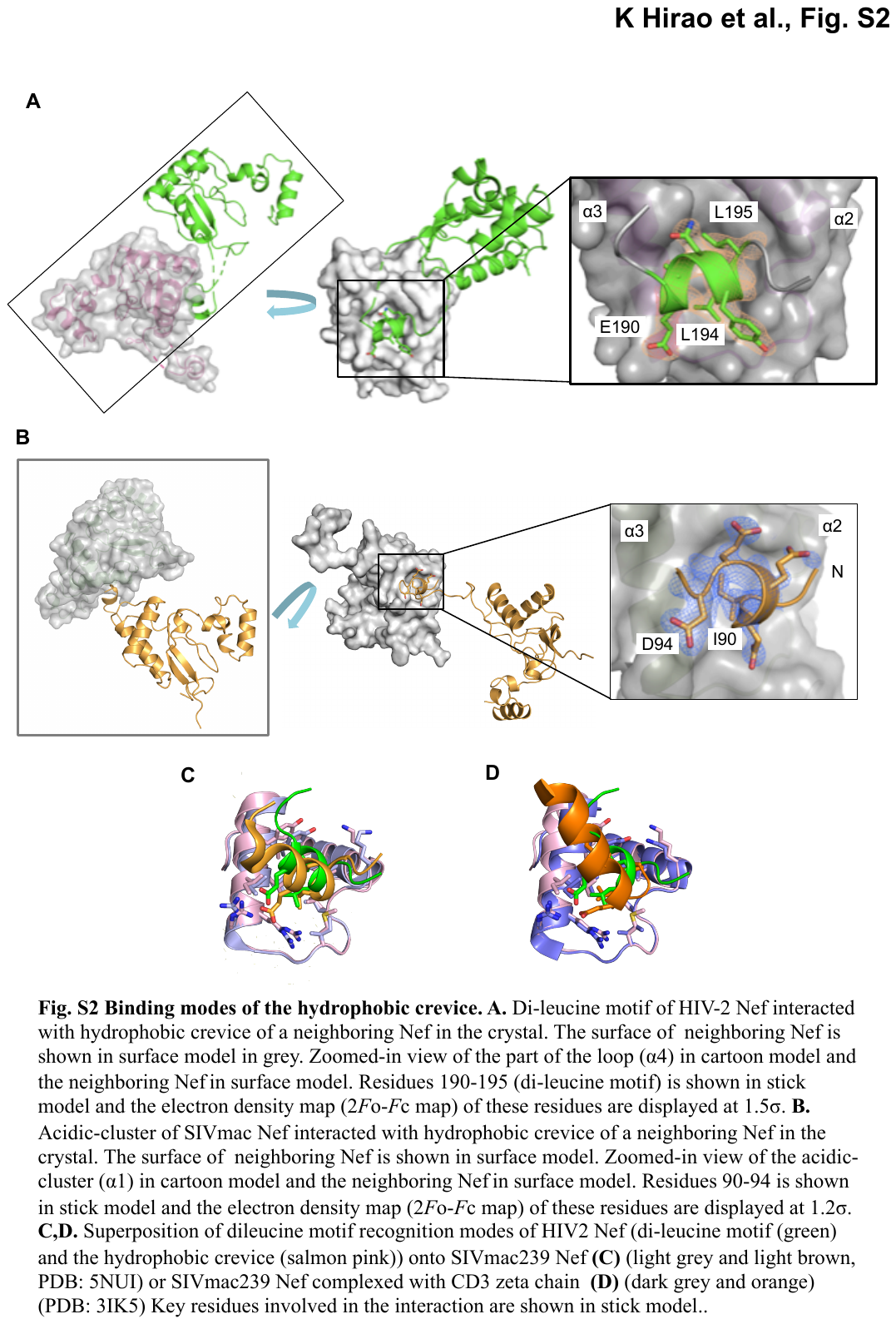

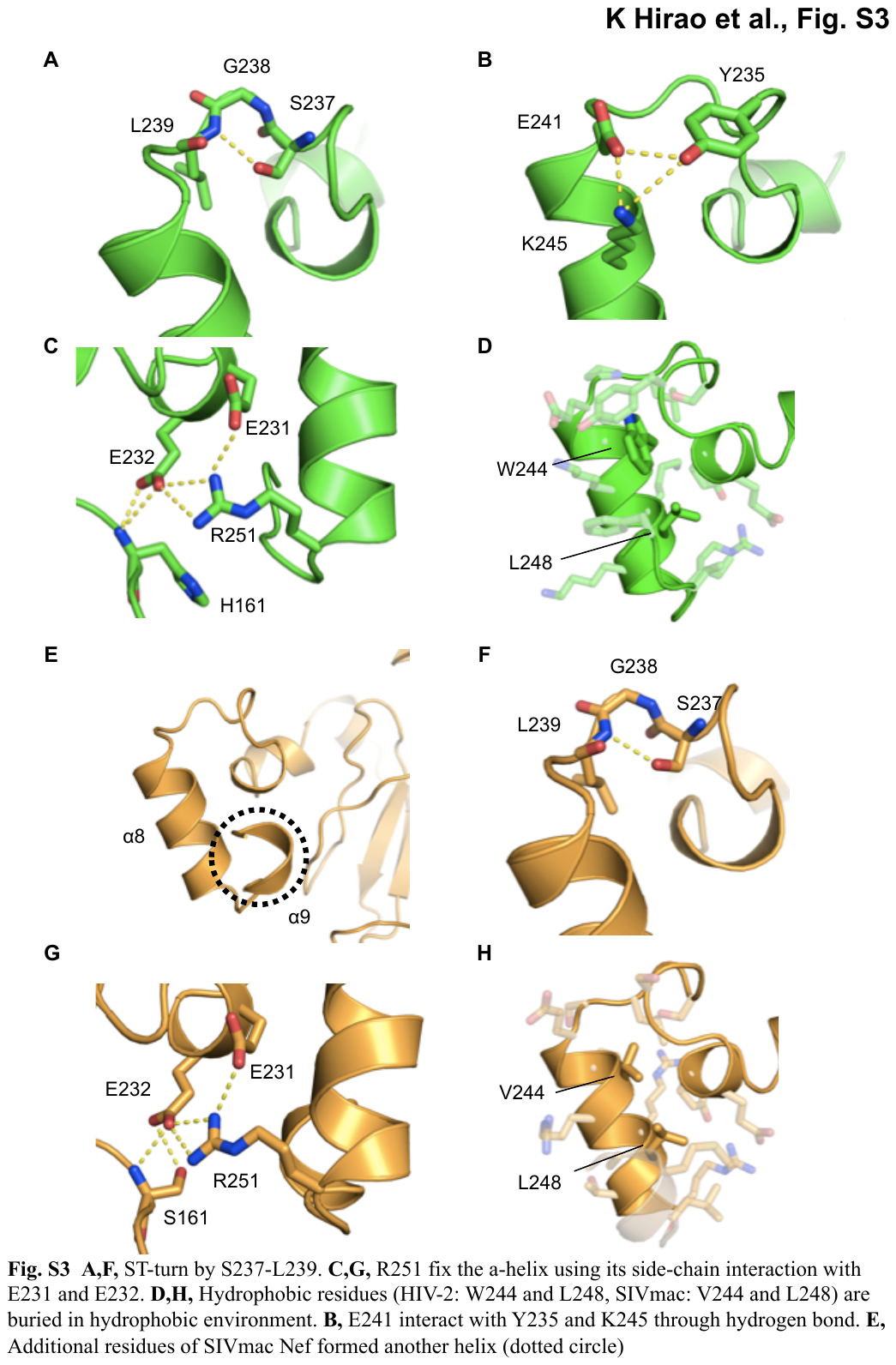

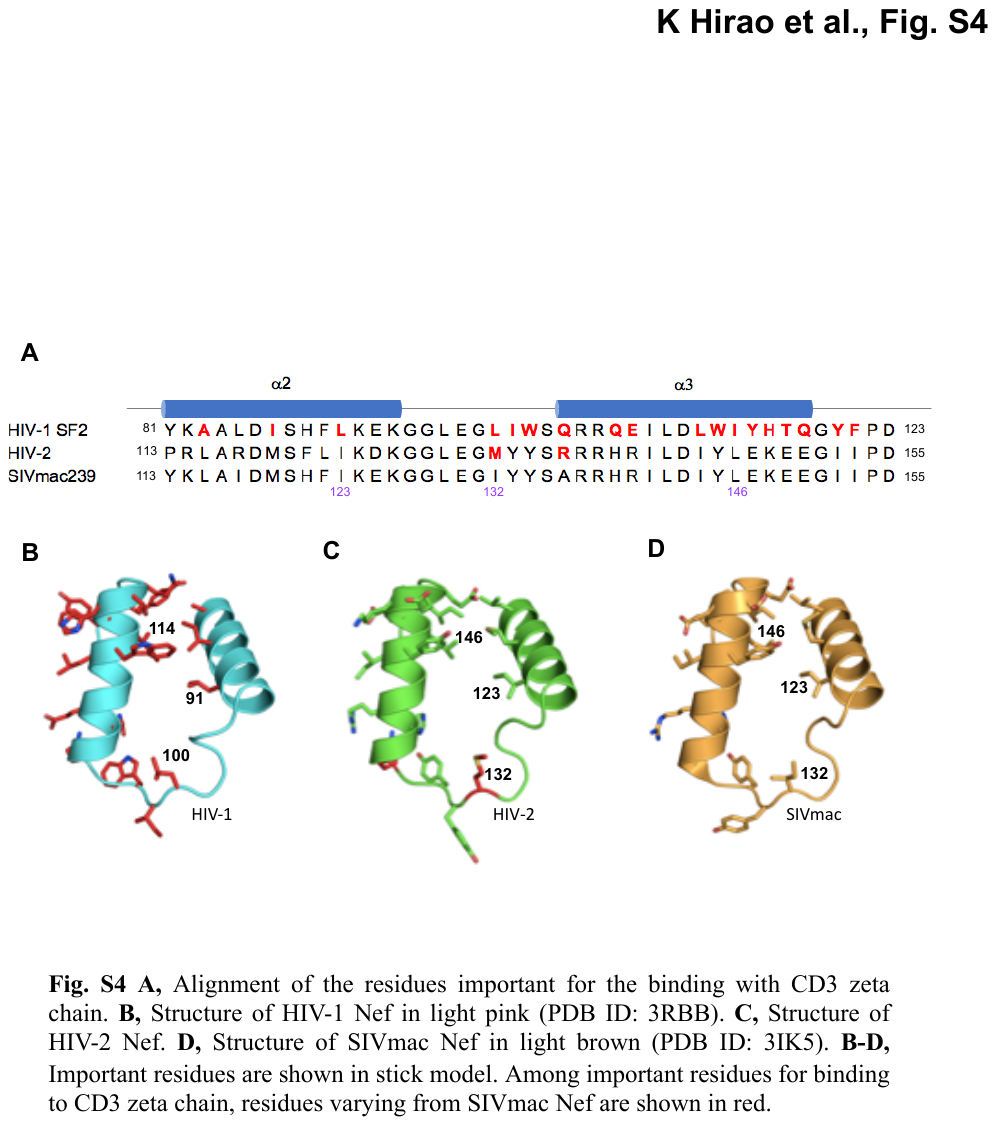

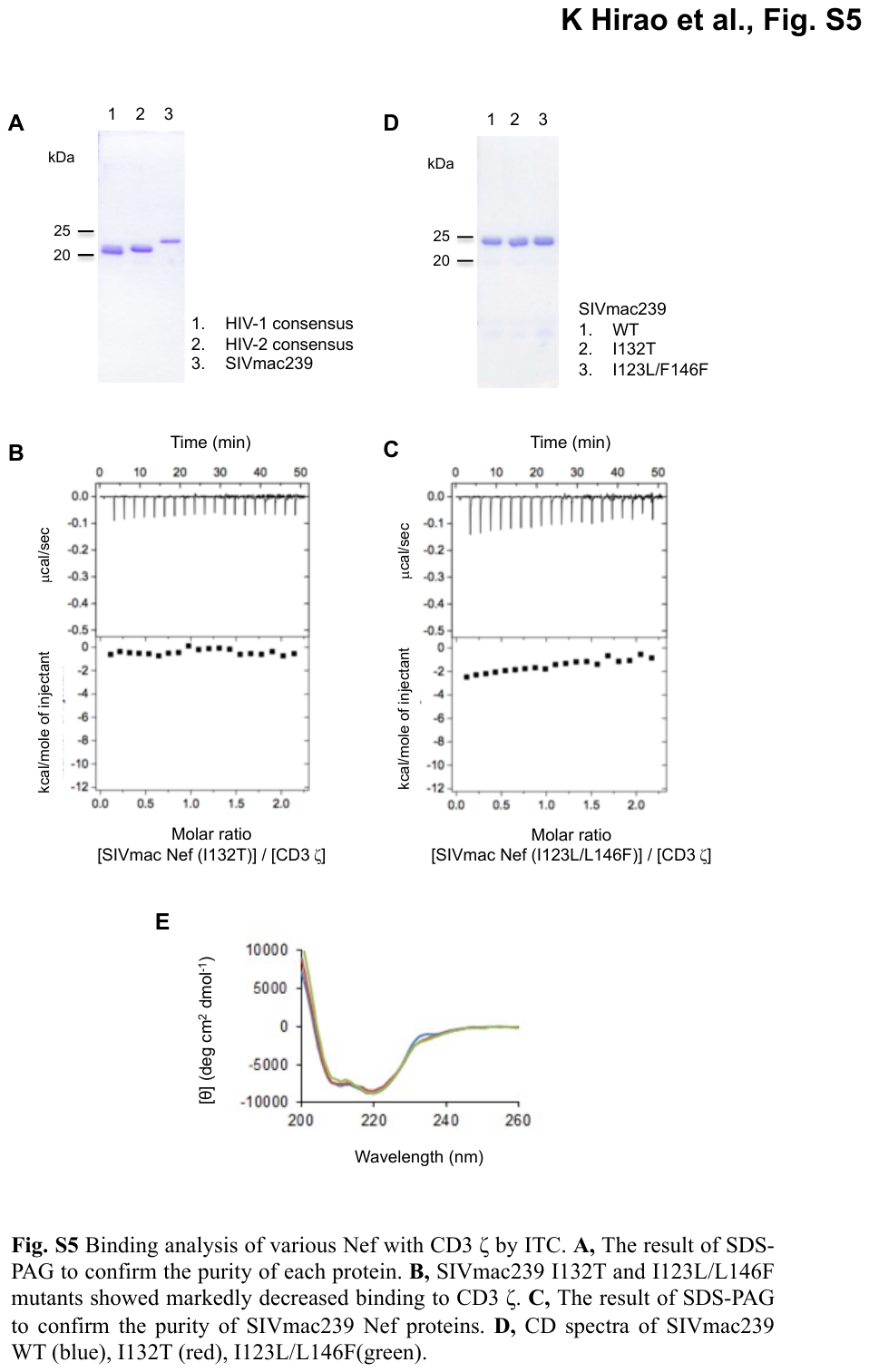

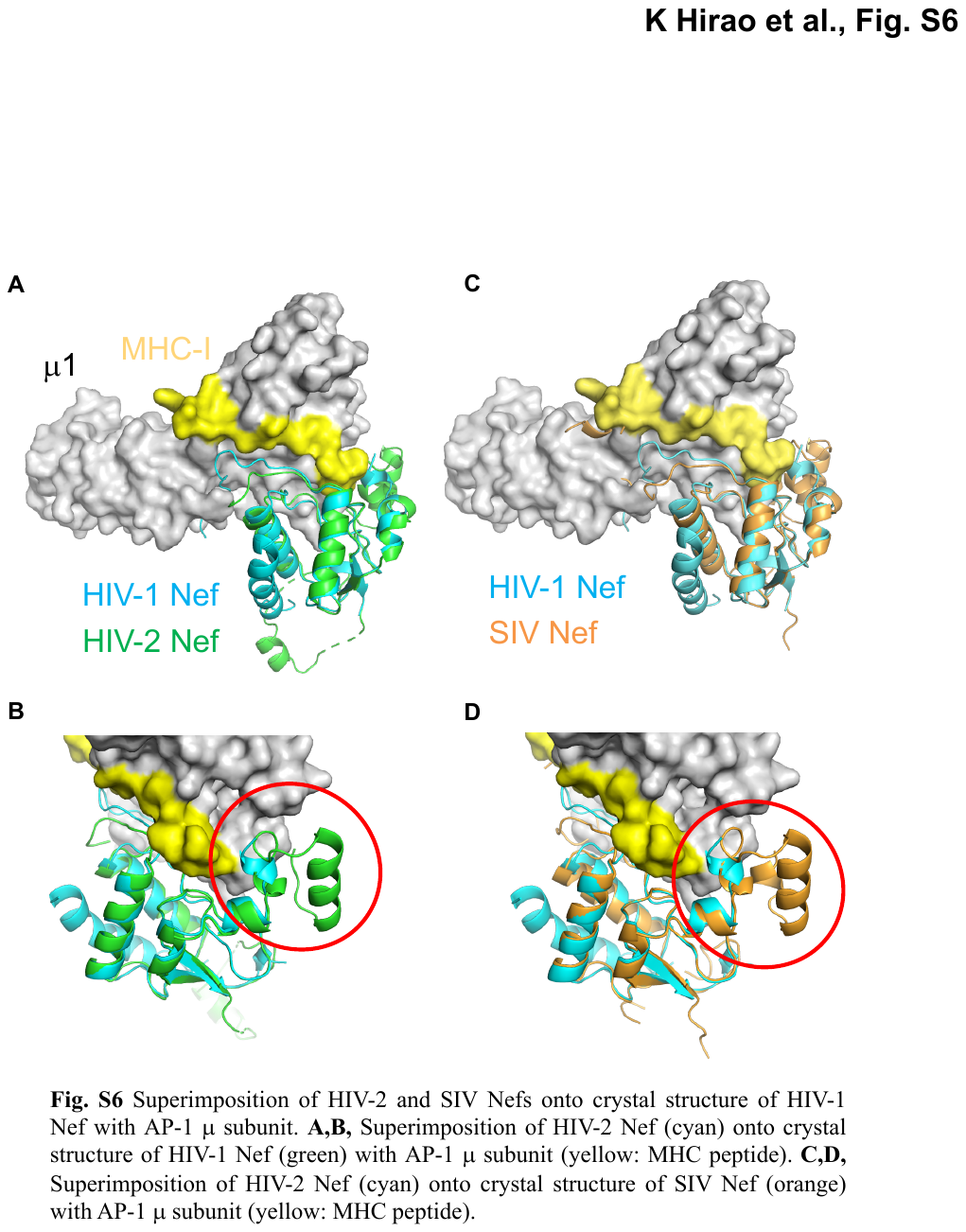
